# Cell therapy with human iPSC-derived cardiomyocyte aggregates leads to efficient engraftment and functional recovery after myocardial infarction in non-human primates

**DOI:** 10.1101/2023.12.31.573775

**Authors:** Ina Gruh, Andreas Martens, Serghei Cebotari, Annette Schrod, Alexandra Haase, Caroline Halloin, Wiebke Triebert, Tobias Goecke, Morsi Arar, Klaus Hoeffler, Paul Frank, Karen Lampe, Amir Moussavi, Veronika Fricke, Nils Kriedemann, Monika Szepes, Kerstin Mätz-Rensing, Jörg Eiringhaus, Anna-Lena de Vries, Ina Barnekow, Claudia Serrano Ferrel, Stephan Hohmann, Merlin Witte, Tim Kohrn, Jana Teske, Victoria Lupanov, Annika Franke, Mark Kühnel, Danny Jonigk, Susann Boretius, Christian Veltmann, David Duncker, Andres Hilfiker, Axel Haverich, Robert Zweigerdt, Ulrich Martin

**Affiliations:** Leibniz Research Laboratories for Biotechnology and Artificial Organs and Dept. of Cardiac, Thoracic, Transplantation and Vascular Surgery, Hannover Medical School, Cluster of Excellence REBIRTH, Hannover, Germany; German Primate Center, Leibniz Institute for Primate Research, Göttingen, Germany; Functional Imaging Laboratory, German Primate Center, Leibniz Institute for Primate Research, Göttingen, Germany; DZHK (German Center for Cardiovascular Research), partner site Göttingen, Germany; Hannover Heart Rhythm Center, Department of Cardiology & Angiology, Hannover Medical School, Germany; Institute of Pathology, Hannover Medical School, Hannover, Germany; Johann-Friedrich-Blumenbach Institute for Zoology and Anthropology, University of Göttingen, Germany

## Abstract

**Background:** Functionally coupled large myocardial grafts and a remarkable improvement of heart function in nonhuman primate models of myocardial infarction have been reported after transplantation of human embryonic stem cell-derived cardiomyocytes at relatively high numbers of up to 10^9^ single cell cardiomyocytes - a dose equivalent to total cell loss after myocardial infarction in ∼10 times larger human hearts. To overcome apparent limitations associated with the application of single cells, this pre-clinical study investigated the injection of cardiomyocyte aggregates instead.

**Methods:** Human iPSC-derived cardiomyocyte aggregates were produced in scalable suspension culture. Intramyocardial injection of the aggregates into cynomolgus monkey hearts was conducted two weeks after myocardial infarction induced by permanent coronary artery ligation. Human cell engraftment was assessed after two weeks or three months; functional analyses included continuous telemetric ECG recording and repeated cardiac MRI assessment in comparison to sham treated animals.

**Results:** Treatment with cell numbers as low as 5 x 10^7^ resulted in efficient structural engraftment. Notably, the degree of heart function recovery *in vivo* seemed to correlate with the contractility of the applied cardiomyocytes tested by parallel experiments *in vitro*. Graft-induced non-life-threatening arrhythmias were transient and decreased considerably during the three months follow-up.

**Conclusions:** Transplantation of human iPSC-derived cardiomyocyte aggregates yielded comparable results to the reported application of higher numbers of single cell cardiomyocytes from human ESC, suggesting that the application of cardiomyocyte aggregates facilitates cell therapy development by reducing cell production costs and clinical risks associated with the administration of relatively high cell numbers.

**Clinical Perspective:** What is new?

- In contrast to previously applied single cells, human iPSC-derived cardiomyocyte aggregates (hiCMAs) were transplanted in a non-human primate (NHP) model of MI, to reduce the required cell dose, promote myocardial retention of the graft, and limit the risks for adverse effects. Such low-dose treatment with almost pure ventricular cardiomyocytes produced under GMP-compliant conditions, resulted in the formation of relative large, structurally integrated human grafts in NHP hearts.
- Transient non-life-threatening arrhythmias associated with intramyocardial cell transplantation decreased considerably during the three months follow-up.
- A remarkable recovery of left ventricular function was observed. This recovery notably correlated with the *in vitro* contractility of transplanted cardiomyocyte batches tested in bioartificial cardiac tissues (BCTs), underlining the relevance of a suitable potency assay.

What are the clinical implications?

- Intra-myocardial injection of hiCMAs is a promising treatment modality for the recovery of contractile function after MI; their advanced production, storage and testing revealed in the study facilitate the clinical translation of hiPSC-based heart repair.
- The need for relatively low numbers of cardiomyocytes produced through advanced protocols for scalable suspension culture reduces production costs of adequate cell batches, thereby increasing treatment availability. *In vitro* testing of the produced cell batches is required to ensure treatment efficacy.
- Clinical hiCMA injection can be considered reasonably safe, however, pharmacological prevention and treatment of arrhythmias is required and temporary implantation of a cardioverter-defibrillator (ICD) could be considered.

## Introduction

Heart failure (HF) remains a major cause of morbidity and mortality worldwide ^1^. Cellular therapies based on human induced pluripotent stem cells (hiPSCs) for the first time offer the *de novo* generation of bona fide human cardiomyocytes (CMs) *in vitro* for replacing lost heart muscle cells *in situ*. Currently, iPSC-based approaches are considered as innovative but complex therapeutic concepts for treating heart failure resulting from conditions such as ischemic cardiomyopathy (ICM) or myocardial infarction (MI). While critical research on safety aspects of iPSC-based cell transplants is ongoing ^2^, substantial progress on iPSC technologies was achieved, including the development of controlled, efficient bioprocesses for the expansion and lineage-directed differentiation of iPSCs into functional well-characterised cardiomyocytes (iPSC-CMs) at clinical scale ^3–5^.

Preclinical studies investigating pluripotent stem cell (PSC)-derived CMs were conducted already in non-human primates (NHP), since the high heart beating rates in rodent models (with the possible exception of guinea pigs ^6^) prevent the proper electromechanical coupling of human CMs ^7, 8^. In addition, the survival of human cells in pig hearts has been limited so far ^9^. Recent NHP studies demonstrated the formation of electrically coupled and well-structured cardiac muscle islands from injected human embryonic stem cell (hESC)-CMs in a xenogeneic approach ^10, 11^, and an equivalent allogeneic study using NHP iPSC-derived CMs ^12^ as well. In that latter work, Shiba et al. reported a significant increase in ejection fraction (EF) and fractional shortening after sub-acute MI and injection of 4 x 10^8^ NHP iPSC-CMs. In a more recent study, the transplantation of 7.5 x 10^8^ single cell-dissociated hESC-CMs into the infarcted heart in a NHP model also suggested the substantial improvement of heart function 3 months after cell treatment ^11^. Notably, however, since the adult human heart is ∼10 to 12 times larger compared to Macaca fascicularis ^13^, these CMs numbers would translate at least into 4 to 7.5 x 10^9^ cells per human patient. The need for such high cell doses creates substantial hurdles for the (commercially viable) cell production and raises additional safety concerns in the clinical setting. The transplantation of PSC-CMs into NHP hearts was also associated with adverse events: ventricular arrhythmias of different characteristics were observed in several cases and persisted throughout the observation period ^10–12^, requesting further investigations before the clinical translation. As of yet, the functional data provided by the NHP studies have to be considered preliminary in view of the small numbers of treated animals and it also remains untested whether the functional heart recovery depends on the contractile properties of the applied CMs or other mechanisms.

Here we report the treatment of cynomolgus monkeys (Macaca fascicularis) with human iPSC-derived cardiomyocytes. In contrast to the application of single cells as in previous NHP studies, we have specifically transplanted cardiomyocyte aggregates (hiCMAs) produced by scalable, chemically defined suspension culture ^4, 5^. This approach aims at testing whether the aggregate delivery, which remains compatible with the administration via intra-myocardial cell injection, promotes the retention, survival and engraftment of transplanted cells and allows to use a substantially lower dose compared to other studies. We demonstrate that the application of well characterized hiCMAs leads to the formation of relatively large grafts despite using relatively low cell numbers, i.e. 5 x 10^7^ CMs per NHP heart. Our data also suggest a remarkable improvement in the left ventricular ejection fraction (LV-EF) compared to infarct-only controls. Notably, a parallel *in vitro* potency assay suggests that higher contractile forces of individual CM preparations do correlate with a functional recovery of hearts treated with the respective cells. Furthermore, we have observed cardiac arrhythmias of different types, frequency and duration after MI induction and subsequent cell transplantation. In analogy to similar studies, these arrhythmias were not life-threatening and gradually decreased over the 3 months observation period.

Together, despite limitations inherent to the NHP model, the study reveals robust proof-of-concept for our hiCMA-based treatment strategy to rescue damaged hearts, thereby providing a realistic option for clinical trials in the near future.

## Methods

### Generation of cardiomyocyte aggregates for transplantation and *in vitro* potency test

Human iPSC cultivation, chemically defined cardiac differentiation, and cell/ aggregate analysis was performed essentially as described ^3, 5^. In brief, the hiPSCs reporter line MHHi0001-A-11 (‘*Amber*’, constitutively expressing Venus(nucmem) ^14^ under control of the CAG promoter from the AAVS1 locus) or MHHi0001-A-5 (‘*Ruby*’, constitutively expressing RedStar(nucmem) and the Calcium sensor GCaMP6f under control of the CAG promoter from the AAVS1 locus ^15^) were used for single cell inoculation and directed cardiac differentiation in suspension culture in 500 mL Disposable Spinner Flask with Vent Cap (Corning), stirred at 60 RPM. After check point 1 (CP1; see Fig. 1A), resulting cardiomyocyte aggregates (hiCMAs) were cultured in RB^+^ medium (RPMI-B27 with insulin, Gibco; plus Penicillin/Streptomycin, Sigma-Aldrich) until check point 2 (CP2; see Fig. 1A). The hiCMA diameter was assessed by microscopic picture analysis and the CM content by flow cytometry. At CP2, hiCMA were collected and used for cell transplantation or for the generation of bioartificial cardiac tissue (BCT) ^5, 16^ to assess CM contractility *in vitro*. For details see supplementary methods.

**Figure 1:**
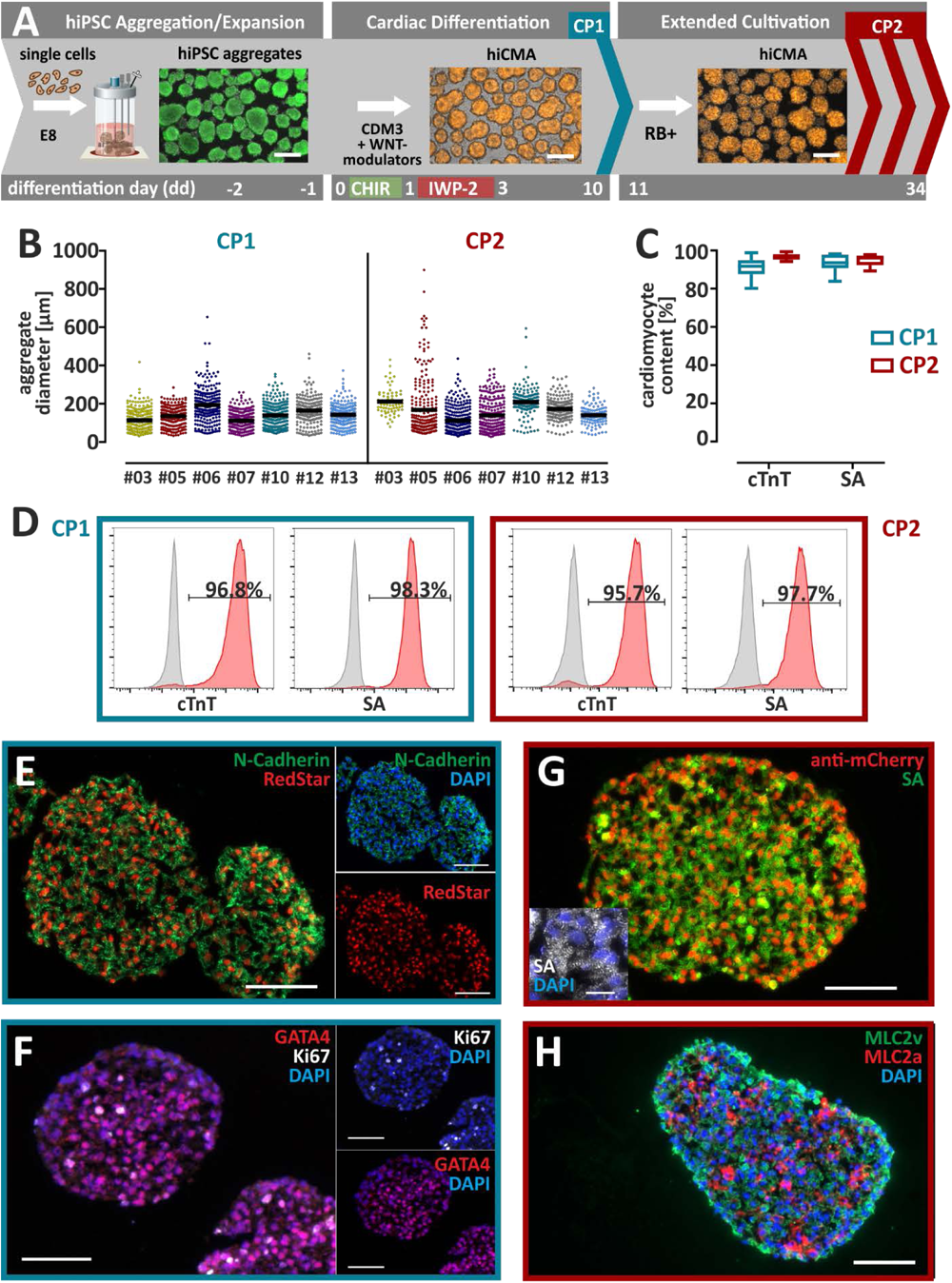
hiPSC-derived enriched cardiac aggregates for transplantation studies can be produced and maintained under defined and scalable culture conditions. **A)** Differentiation workflow. Inoculation of a hiPS cell suspension in E8 Medium in stirred spinner flasks resulted in the formation of cell aggregates within 2 days (fluorescence microscopy (FM) picture of calcein viability stain). For directed cardiac differentiation from dd0 onwards, WNT-modulators (CHIR and IWP-2) are supplemented in Cardiac Differentiation Medium 3 (CDM3). From dd11 onwards, CDM3 is replaced by RB+ Medium for extended hiCMA cultivation until transplantation on CP2. On process Check Point 1 (CP1; end of differentiation) and CP2 (dd18-dd34, time point of tx) samples of cardiomyocyte aggregates were harvested for analysis (hiCMAs; FM picture of RedStar expression in CAG-RedStar(nucmem)-AAVS1 reporter hiCMAs from hiPSC MHHi0001-A-5 ‘*Ruby*’; Scale bars: 100 µm). **B)** Independently differentiated hiCMA batches used for transplantation experiments displayed similar aggregate diameters assessed on CP1 and CP2, respectively. Each dot represents an individual aggregate; bars represent the resulting median diameter; #03 - #07 *Amber* hiCMAs; #10 - #13 *Ruby* hiCMAs. **C)** Flow cytometry (FC) for independent differentiation runs used for transplantation experiments (n=4-6) indicated a typical CM content of ∼95-98% after staining of dissociated hiCMAs on CP1 and CP2, respectively, for cardiac troponin T (cTnT) and α-sarcomeric actinin (SA). **D)** Representative FC data show high levels of cardiac marker expression (red) compared to controls (grey)**. E-H)** Immunofluorescence (IF) analysis of cryosections of representative hiCMAs from CP1 (in E&F) and CP2 (in G&H) for expression of N-cadherin, the cardiac marker α-sarcomeric actinin (SA) and the transgene RedStar (unstained in E; stained for better visibility with a cross-reacting anti-mCherry mAb in **G**). **F)** IF at CP1 revealed expression of the cardiac progenitor marker GATA4, as well as the cell cycle marker Ki-67 (Ki67). **H)** Cryosections of representative hiCMAs from CP2 show expression of both ventricular and atrial myosin light chain (MLC2v and MLC2a). Scale bars: 100 µm; in the inset in G: 20 µm.

### Animal Care

Cynomolgus monkeys (Macaca fascicularis; 1 female, 6.8 kg; 14 male 7.3 - 12.8 kg) were kept at the German Primate Center under conditions according to §§7-9 of the German Animal Welfare Act, adhering to the European Union guidelines (EU directive 2010/63/EU) on the use of non-human primates for biomedical research. The experiments were approved by the Lower Saxony State Office for Consumer Protection and Food Safety. To prevent early infarction-related arrhythmias, animals were treated with amiodarone (100 mg day^-1^, oral) for 14 days starting 5 days before induction of the myocardial infarction (MI). Metoprolol was administered twice daily (50 mg) for 11 days starting 2 days before surgery. Immunosuppression started 5 days before cell transplantation with daily injections of cyclosporine A (10 mg kg^-1^ i.m. for 15 days, followed by a maximum of 7.5 mg kg^-1^ or an adjusted dose to maintain serum trough levels of 600 µg/L) and methylprednisolone (2.5 mg kg^-1^ i.m.). In addition, belatacept (CTLA4-Ig, 10 mg kg^-1^ i.v.), a T-cell costimulatory signal blocker suitable for cynomolgus monkeys ^17^ was administered on the day of cell transplantation, and on day 4, 14, 28, and 56 after cell transplantation. Marbofloxazin (4 mg kg^-1^ i.m. during surgery) and amoxicillin (0.05 mg kg^-1^; s.c.) were administered to prevent opportunistic infections. Post-surgery, wounds were carefully examined and animals were regularly monitored by laboratory (for hematocrit, hemoglobin levels) and Primate Center staff (for signs of distress indicating post-procedure pain). Finally, all animals were euthanized under deep anesthesia with pentobarbital after administration of 500IE/kg heparin i.v.. Tissue processing, histology, immunofluorescence staining, and image analyses were performed as described in the supplementary methods; primary and secondary antibodies are listed in tables S1, S2 and S3.

### Animal model of myocardial infarction and cell transplantation

Animals underwent left anterior thoracotomy and permanent ligation of the left anterior descending artery (LAD) distal to the first diagonal branch for the induction of left ventricular MI under general anesthesia. MI was confirmed by a pale ischemic area and by serum assays for cardiac troponin (cTnT) and creatine kinase (CK and CK-MB) 1 day after surgery. Trans-esophageal cardiac echocardiography was performed before and after MI. Two weeks after MI, a re-thoracotomy was performed under general anesthesia. The heart was exposed and hiCMAs in phosphate-buffered saline (PBS) were injected into the peri-infarct region of the left ventricular wall via 10-12 injections of 50 µl volume each with a 26-G needle. Each hiCMA sample injected per heart contained ∼50 million CMs (∼70 million CMs for animal #01, only) calculated based on the analysis of parallel, representative hiCMA aliquots dissociated into single cells for counting). An equal volume of phosphate buffered saline (PBS) was injected in control animals. For details see supplementary methods.

### ECG monitoring

Continuous monitoring of clinically relevant arrhythmias was established using an implantable telemetry device (PhysioTel Digital M01, Data Sciences International (DSI), St. Paul, MN, USA). The device was implanted at the time of LAD ligation surgery during general anesthesia. DSI Ponemah® version 6.20 with Pattern Recognition Option (DSI, St. Paul, MN, USA) was used for ECG telemetry data collection and analysis. For details see supplementary methods.

### Cardiac MRI

The anesthetized and intubated animals underwent MRI measurements before MI (pre-MI), 2 weeks after MI (directly before cell transplantation), and 2 and 12 weeks after cell transplantation with a 3T MR-system (Magnetom Prisma, Siemens Healthineers, Erlangen, Germany) using a 16-channel multipurpose coil (Variety, Noras MRI products, Hoechberg, Germany). Left ventricular function was assessed using an ECG-gated segmented cine Fast Low-Angle Shot (FLASH) technique or free breathing real-time MRI in case of corrupted ECG signal ^18^. The standard volumetric and functional parameters of the left ventricle (end-diastolic volume (EDV), end-systolic volume (ESV), stroke volume (SV), and ejection fraction (EF)) were assessed. Due to variance of pre-MI LV-EF, the relative LV-EF (LV-rEF) was defined as the LV-EF normalized by the pre-MI EF of the same animal. Despite systematic differences in the estimated absolute values of EF acquired by cine and Real-Time MRI, no significant differences in rEF are expected based on our previous work extensively comparing the two methods for their applications in macaques ^18^. Maps of the T1-relaxation time were acquired before and after contrast agent administration (Gadobutrol/Gadovist^TM^, 0.1 mmol/kg) using a fast Real-Time single-shot inversion recovery FLASH technique ^19^. For details see supplementary methods.

### Statistical analysis

The GraphPad Prism Version 9 (GraphPad Software, Inc., La Jolla, CA, USA) was used for statistical analyses. Unless indicated otherwise, descriptive statistical analysis was expressed as Mean ± Standard Deviation. Continuous numerical variables were compared using the student’s t-test for independent samples, significance was evaluated by two-tailed testing, and assumed at p<0.05. To compare more than two groups, a two-way ANOVA with Tukey’s or Bonferroni’s multiple comparisons test was performed. A *p*-value <0.05 was considered statistically significant. For the inter-observer comparison, the intraclass correlation coefficient (ICC) was calculated using repeated measures ANOVA ^20^ and a Bland-Altman plot to graph the data ^21^.

## Results

### Cardiomyocyte purity is maintained after prolonged aggregate suspension culture

We have recently established advanced cardiomyogenic differentiation of hPSC in chemically defined suspension culture, enabling the robust induction of ventricular-like cardiomyocytes (CMs) at ∼95-98% purity for numerous cell lines ^5^. Here, this protocol was extended beyond process assessment on “checkpoint 1” (CP1, d10) by additional cultivation of the resulting cardiac aggregates (hiCMA) in stirred suspension culture for further 9-24 days until “checkpoint 2” (CP2; dd19-dd34) (Fig. 1A, Table1).

Despite some inter-experimental variability in the average diameter of hiCMAs (which was around ∼200 µm) and the aggregate size distribution (Fig. 1B), no apparent differences between CP1 versus CP2 samples were observed. Flow cytometry (FC) specific to sarcomeric markers actinin (SA), and cardiac troponin T (cTnT) revealed stable CM content (typically >95%) across independent process runs (Fig. 1C) with equal expression levels for cTnT and SA at CP1 and CP2 (Fig. 1D). Notably, hiCMAs derived from both hiPSC-reporter cell lines used in this study, *Amber* (for animals #03 - #07) and *Ruby* (for animal #10, #12 & #13) ^15^, showed similar aggregate sizes, CM content and markers expression.

Typical hiCMAs were round shaped and showed the expected nuclear localization of the reporter transgene (Fig. 1E). Immunostaining of sections at CP1 revealed early cardiac marker GATA4 and proliferation marker Ki67 expression, suggesting an expected ‘fetal-like’ status (Fig. 1F). At CP2, SA^pos^ CMs were homogeneously distributed without particular cell alignment, but with a cross-striated pattern showing sarcomeric structures (Fig. 1G, see inset); persistent RedStar reporter gene expression was detected with an anti-mCherry antibody (Fig. 1G). The aggregates contained mostly MLV2v^pos^ and a portion of MLC2v^pos^/MLC2a^pos^ cells (Fig. 1H). hiCMAs showed a contractile phenotype *in vitro* (see video 1 for *Amber* derivatives and video 2 showing the calcium transient-indicating flashing of *Ruby*-derived hiCMAs). At CP2, hiCMAs were directly used - without any dissociation or other downstream processing steps - for transplantation into the heart in our non-human primate (NHP) model of myocardial infarction (MI) and for an *in vitro* potency assay.

### *In vitro* potency assessment reveals cell line-dependent differences in contractile force development of hiCMA-derived CMs

Parallel to the cell transplantation *in vivo*, individual hiCMA batches from CP2 were tested for contractile bioartificial cardiac tissue (BCTs ^16^) formation *in vitro*. CMs from both reporter cell lines formed spontaneously contracting tissues (termed *Amber*-BCTs and *Ruby*-BCTs, respectively), which were analyzed 14 days after tissue production, reflecting CP3 of the *in vivo* study (Fig. S1A). BCTs contained aligned SA^pos^ cardiomyocytes (Fig. S1B) and showed typical morphological changes, i.e. tissue thinning, due to matrix remodeling over time (Fig. S1C,E). Spontaneous contraction forces were measured to assess function (example in Fig. S1D); for *Ruby*-BCTs expressing GCaMP6f, video-optical assessment of calcium oscillations showed synchronized tissue contractions (Fig. S1F, G). Interestingly, active forces were generally higher in *Amber*-BCTs (3.2 ± 1.3 mN vs. 0.7 ± 0.6 mN in *Ruby*-BCTs; Mean ± SD for n=8-17 BCTs from 3 experiments each; p<0.05 for a two-tailed nested t-test; Fig S1H), passive forces indicating tissue stiffness were higher in *Ruby*-BCTs with values up to 14.2 ± 5.0 mN (Mean ± SD, n=12 BCTs; Fig. S1I).

### Infarction-related loss of myocardial function in the NHP model can be correlated with serum levels of troponin T as a reliable biomarker

An MI model in cynomolgus monkeys (n=10) was established by permanent coronary artery ligation under general anesthesia (Fig. 2A). Directly following the ligation, the affected territory appeared pale and intra-operative trans-oesophageal echocardiography showed dyskinetic myocardial regions (data not shown). Untreated MI led to extensive myocardial scar formation after 3 months (Fig. 2A).

**Figure 2:**
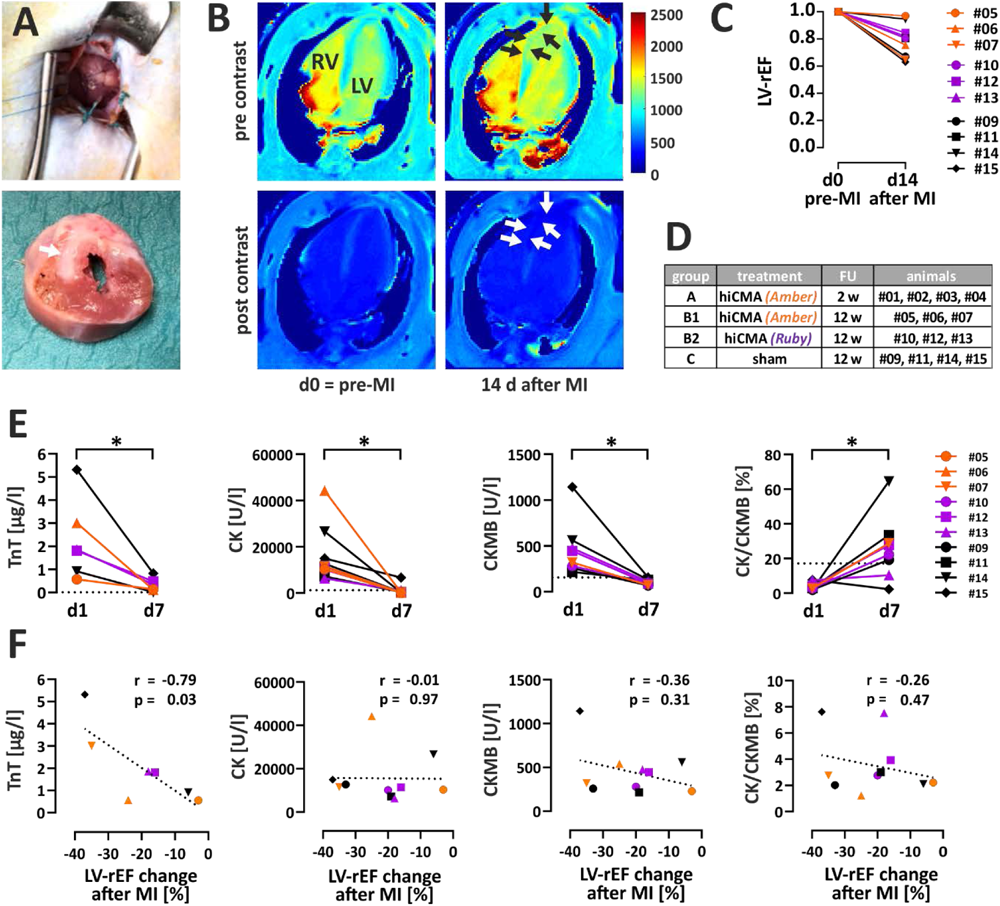
Induction of myocardial infarction in cynomolgus monkeys. **A)** Myocardial infarction was induced by permanent coronary artery ligation with a 6-0 prolene suture after exposing the heart via left lateral thoracotomy under general anaesthesia. Lower panel: Macroscopic assessment of scar formation after explantation in a short-axis heart slice of an untreated animal (#09) 3 months after MI. White arrow indicates extensive scar spanning septum and left-ventricular wall. **B)** Exemplary cardiac MR images of the healthy animal #09 myocardium before (pre-MI) and the infarcted myocardium 14 d after myocardial infarction confirm local changes in tissue composition. Pre-contrast (above) and post-contrast (below) maps of T1-relaxation time [ms] (see color scale) clearly visualize the fibrotic scar region (white arrows) in the septum and the apex of the myocardium. **C)** Cardiac MRI was used to quantify changes in relative left ventricular ejection fraction (LV-rEF) on d14 after MI compared to pre-MI values (normalized to 1); while overall reduction of LV-rEF after MI was significant (*, p < 0.05 for paired t-test with n=10 animals), very small changes were observed in animals #05 and #14 (see table 1 for individual data). **D)** Infarcted animals were assigned to the treatment groups A and B1 (treatment with hiCMA ‘*Amber’*), group B2 (treatment with hiCMA ‘*Ruby’*), and group C (sham, black). **E)** Serum levels of clinically used infarction markers troponin T (TnT), creatine kinase (CK), creatine kinase MB isoenzyme (CK-MB), and their ratio were determined 1 and 7 days after coronary artery ligation in animal groups B1, B2, and C (n=7-10; based on blood sample and/or assay availability). Dotted lines represent mean levels measured in healthy control animals unrelated to this study (n=6). **F)** Correlation analyses of serum levels of infarction markers TnT, CK, CK-MB, and CK-MB/CK on d1 after coronary artery ligation and functional impairment of cardiac contractility on day 14 assessed by MRI (LV-rEF change after MI [%] = (LV-rEF d14 – LV-rEF d0)*100) for groups B1, B2, and C (Pearson correlation coefficient (r) and p value for two-tailed analysis with n=7-10).

**Table 1.** Data for individual macaques in MRI study Figures 1-6.

|  |  |  |  | Pre-infarction |  |  | Post-infarction baseline |  |  | 2 weeks after treatment |  |  | 12 weeks after treatment |  |  |
| --- | --- | --- | --- | --- | --- | --- | --- | --- | --- | --- | --- | --- | --- | --- | --- |
| Anima I # | Sex | Age (years) | Weight (kg) | LVED V (ml) | LVESV (ml) | LVEF (%) | LVED V (ml) | LVESV (ml) | LVEF (%) | LVED V (ml) | LVESV (ml) | LVEF (%) | LVED V (ml) | LVESV (ml) | LVEF (%) |
| **Group B1 - hiCMA iPSC1 transplantation** |  |  |  |  |  |  |  |  |  |  |  |  |  |  |  |
| #05* | M | 6.1 | 11.7 | 11.7 | 6.8 | 41.7 | 12.4 | 7.4 | 40.6 | 10.1 | 5.9 | 41.6 | 11.6 | 6.6 | 43.0 |
| #06 | M | 7.9 | 10.5 | 15.1 | 5.3 | 64.8 | 16.0 | 8.2 | 49.0 | 14.1 | 5.4 | 61.6 | 15.6 | 6.6 | 57.8 |
| #07 | M | 7.5 | 8.8 | 15.6 | 5.6 | 64.3 | 16.6 | 9.7 | 41.6 | 12.3 | 7.1 | 42.4 | 13.4 | 6.2 | 54.1 |
| **Group B2 - hiCMA iPSC2 transplantation** |  |  |  |  |  |  |  |  |  |  |  |  |  |  |  |
| #10* | M | 7.3 | 11.8 | 9.8 | 5.6 | 42.9 | 8.3 | 5.5 | 33.6 | 8.7 | 5.5 | 37.1 | 7.8 | 5.1 | 36.0 |
| #12 | M | 7.3 | 12.3 | 12.0 | 4.5 | 64.0 | 12.7 | 5.9 | 54.9 | 12.3 | 6.0 | 52.0 | 14.5 | 6.6 | 55.3 |
| #13 | M | 7.3 | 9.0 | 16.4 | 6.6 | 60.8 | 15.3 | 7.8 | 49.5 | 14.2 | 6.8 | 51.1 | 16.3 | 8.4 | 48.7 |
| **Group C - Sham control** |  |  |  |  |  |  |  |  |  |  |  |  |  |  |  |
| #09 | M | 7.2 | 9.8 | 17.8 | 7.7 | 56.8 | 20.1 | 12.4 | 38.1 | 18.2 | 11.3 | 37.8 | 19.3 | 11.5 | 40.5 |
| #11 | M | 3.7 | 7.8 | 10.9 | 3.5 | 67.5 | 10.6 | 4.9 | 54.1 | 9.8 | 4.0 | 59.3 | 11.0 | 5.3 | 51.6 |
| #14 | M | 7.7 | 10.5 | 12.3 | 4.5 | 63.7 | 15.1 | 6.0 | 60.1 | 15.4 | 5.4 | 64.8 | 14.2 | 5.8 | 59.5 |
| #15 | M | 7.6 | 9.3 | 13.9 | 6.0 | 56.7 | 15.4 | 9.9 | 35.7 | 16.3 | 10.1 | 38.2 | 18.7 | 11.1 | 40.8 |

MRI data are given as Mean from two independent observers; LVEDV. Left ventricular end-diastolic volume; LVESV. Left ventricular end-systolic volume; LVEF. Left ventricular ejection fraction; \* lower absolute values for MRI parameters due to analysis with real time mode

The volumetric and functional parameters of the left ventricle are summarized in Table1, showing a good inter-observer agreement exemplified for the LV-EF (intraclass correlation coefficient of 0.89, Fig S2B). On d14 after MI, loss of contractile myocardium and late gadolinium enhancement in left ventricular and septal areas near the apex were observed (Fig. 2B and Fig. S2C) in all animals, except animal #05 and #14, which were excluded from further MRI analysis.

Cardiac MRI reveals a significant increase in LV-ESV after MI (7.8 ± 2.4 ml vs. 5.6 ± 1.2 ml before MI, paired t-test, p<0.05 for n=10), while the LV-EDV did not change significantly (paired t-test, p=0.14 for n=10; Fig. S2A). Consequently, LV-rEF was reduced significantly to 0.79 ± 0.12 (Fig. 2C, paired t-test, p<0.05 for n=10). Following MI, individual animals were treated either by intra-myocardial injection of hiCMAs in groups A, B1, and B2, for the two cell lines and time points of analyses, or of DPBS for the sham control group C (Fig. 2D). Assignment to the treatment groups was performed upon sequential inclusion of the available animals into the study, aiming at the best possible age and weight distribution for individual groups (see table 1). On d1 after ligation, clinically used infarction biomarkers troponin T (TnT), creatine kinase (CK), and creatine kinase MB isoenzyme (CK-MB) showed significantly elevated levels that returned back to normal on d7 (Fig. 2E). Only d1 levels of TnT significantly correlated with changes in LV-rEF observed by MRI 14 days after MI (Fig. 2F). Thus, troponin T seems to represent a reliable biomarker for the extent of infarction-related loss of contractile function in our animal model.

### Human iCMAs form large grafts structurally integrated into the primate myocardium

Intra-myocardial injection of hiCMAs was performed after re-thoracotomy under general anesthesia 14 days after MI, reflecting a sub-acute disease state; immunosuppression was initiated 5 days before cell administration (Fig. S3A). A first set of experiments with hiCMAs from *Amber-*iPSCs (group A, n=4) focused on short-term engraftment 2 weeks after hiCMA injection (i.e. CP3, Fig. 3A and S3B). In 4 out of 4 NHP hearts, donor cells were detected: human CMs were identified by staining with an anti-human cTnI antibody not cross reacting with cynomolgus cTnI, as confirmed by exclusive co-staining of the graft’s Venus reporter by an anti-GFP antibody. Grafts of different dimensions were observed, including small, hiCMA-sized areas, which showed structural integration between the host heart muscle fibers, suggesting that individual aggregates do not disintegrate during the injection procedure (Fig. 3B). Relatively large graft areas were detected as well, indicating substantial cell survival (Fig. 3C). The graft size was far beyond the size of individual hiCMAs, strongly suggesting that hiCMAs connected and ‘fused’ *in vivo* to form a tissue. Already on day 14 after transplantation, human cell alignment was visible within the grafts, including clear cross-striation of the donor CMs (Fig. 3D); both aligned and less organized areas were observed in the center of larger grafts. Human cells expressed the CM markers titin, cTnI, SA (Fig. 3B-D, G-I) and cTnT (not shown). The vast majority of grafted human cells solely expressed the ventricular marker MLC2v; only a minority expressed MLC2a (Fig. 3E).

**Figure 3:**
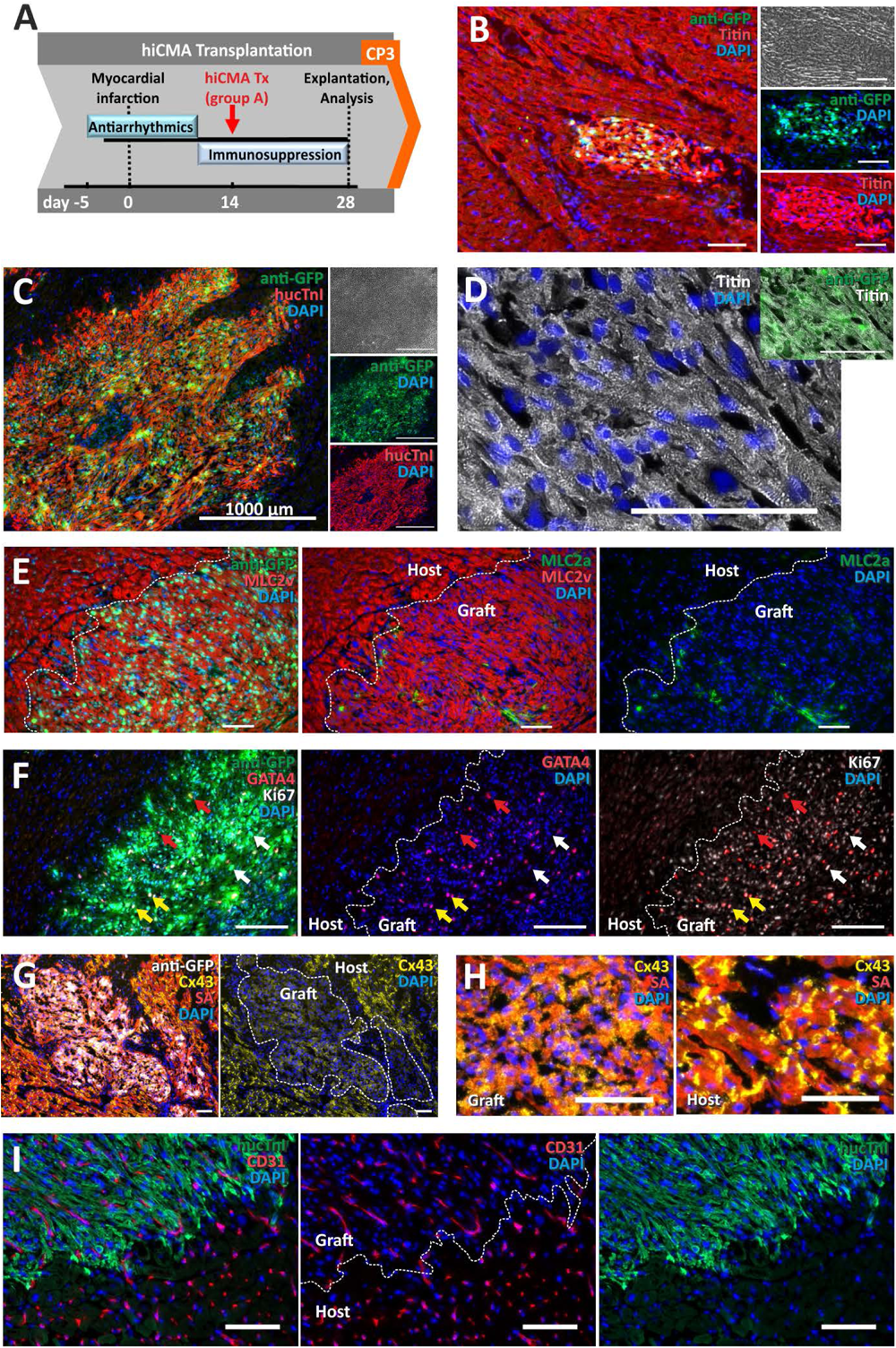
Human iPS-derived cardiomyocyte aggregates (hiCMAs) integrate into the infarcted non-human primate heart and form large matured grafts within 14 days. **A)** Experimental outline of short-term group A with hiCMA transplantation 14 d after MI and heart tissue sampling and analyses after another 14 d on d28 (CP3). **B)** Small human cardiomyocyte (CM) graft presumably originating from a single hiCMA, identified by IF in the host myocardium with an anti GFP antibody (staining the yellow GFP variant Venus, green) and an antibody against Titin (red). **C)** IF also identified large grafts of human CMs in the host myocardium with a species-specific antibody against human cTnI (red) and GFP (staining Venus, green). **D)** Venus(nucmem)^pos^ hiCMA grafts (green) expressing the cardiac marker titin (white) contain aligned, elongated cells with a cross-striated staining pattern that reflects sarcomeric organization. **E)** Characterization of CM subtype marker expression by IF with shows predominantly MLC2v^pos^ CMs (red) in the graft area identified with the anti-GFP mAb (green, left panel), boundary marked with a dotted line. Middle and right panels: Co-staining for MLC2v^pos^ CMs (red) and MLC2a^pos^ CMs (displayed in green). **F)** In human grafts, identified with the anti-GFP antibody (left panel) and marked with a dotted line (middle and right), IF detected GATA^pos^ CMs (red, red arrows), Ki67^pos^ cells (white, white arrows) and GATA^pos^/Ki67^pos^ CMs (yellow arrows). **G)** Connexin 43 (CX43, yellow) is expressed in human grafts but at a lower level than in the surrounding NHP tissue. **H)** CX43 is less organized in human grafts (left) than in the host myocardium (right). **I)** In human grafts, identified with the antibody against human cTnI (green), IF detected CD31^pos^ endothelial cells (red). All nuclei were stained with DAPI (blue); Scale bars 100 µm in B; 1000 µm in C; 100 µm in D-I.

The number of Ki67^pos^ cells was higher in the graft region compared to the surrounding host myocardium (Fig. 3F) including a subset of GATA4^pos^/Ki67^pos^ human CMs, indicating maintenance of an immature embryonic-like expression profile. Expression of connexin 43 (CX43) was significantly lower and less organized in the grafts compared to the host myocardium (Fig. 3G, H). Notably, within the graft regions, also CD31^pos^ endothelial cells (ECs) were abundant (Fig. 3I).

Long-term survival and integration of hiCMAs after MI (group B animals) was explored after 3 months (Fig. 4A) for both *Amber-*hiCMAs (group B1, n=3) and *Ruby-*hiCMAs (group B2, n=3). On d96 after transplantation, human cells were detected in 6 out of 6 NHP hearts in this xenogeneic model. Accompanied by smaller grafts, very large islands of engrafted human CMs spanning more than 5 mm of the host myocardium were detected (Fig. 4B). Depending on the muscle fiber orientation in the analyzed tissue, linear arrangement of CMs with distinct cross-striations was observed (Fig. 4C). The number of MLC2a^pos^ human CMs was further reduced after 3 months, indicating progressive *in vivo* maturation of the predominantly MLC2v^pos^ cells (Fig. 4D&E). Anti CX43 staining suggested cell-cell coupling of individual host and graft cardiomyocytes via gap junctions (Fig. 4F). Moreover, CD31^pos^ cells showed a host-derived vascularization of the newly formed human myocardium (Fig. 4G). No apparent differences were observed for graft integration of the two reporter cell lines *Amber* (in Fig. 4B-D) and *Ruby* (Fig. 4 F,G).

**Figure 4:**
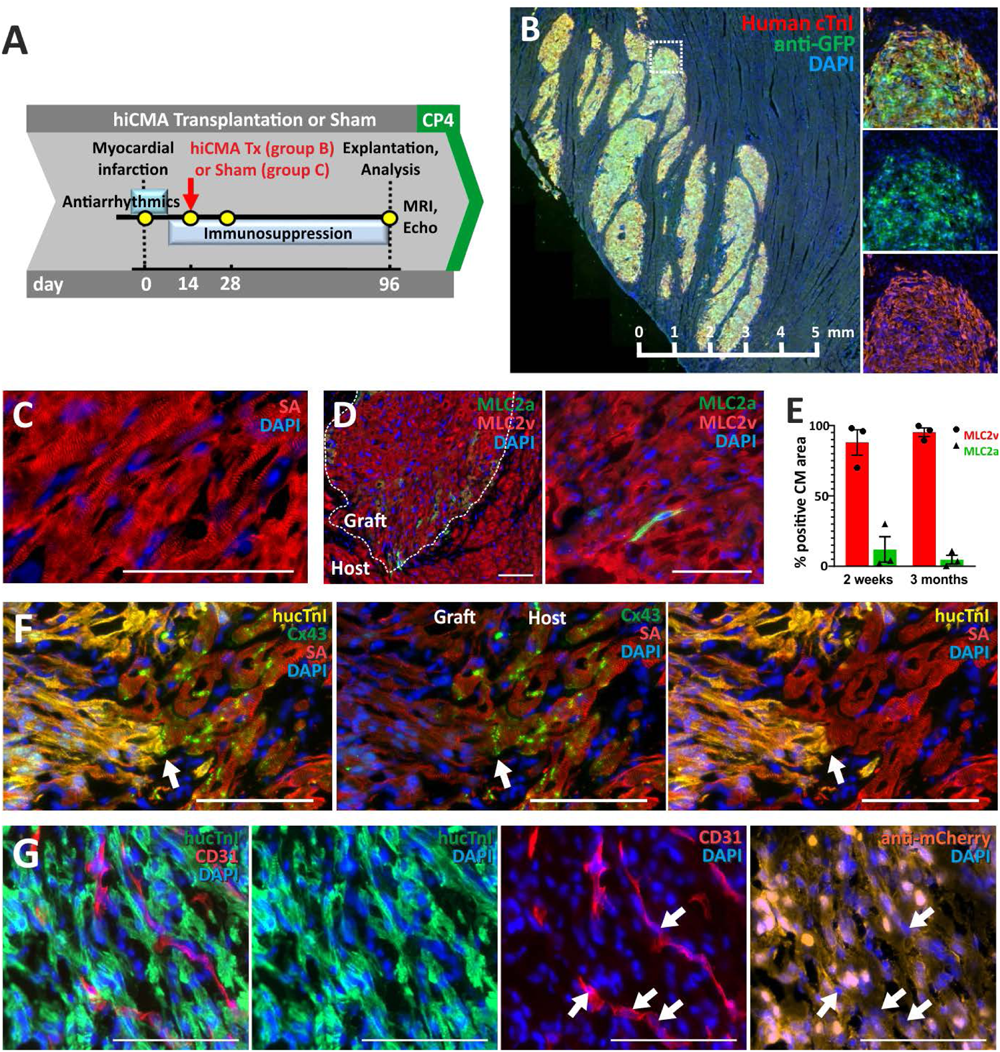
Three months after injection human iCMAs form large grafts structurally integrated into the infarcted non-human primate heart. **A)** Experimental outline of long-term group with hiCMA transplantation 14 d after MI and heart tissue sampling and analyses on day 96 (3 months after cell transplantation, CP4). **B)** Extensive grafts of human CMs in the host myocardium immunostained for human-specific cTNI (red) and the yellow GFP variant Venus(nucmem) (green). **C)** Anti sarcomeric actinin (SA) staining shows aligned human graft myocardium with distinct cross-striation. **D)** IF for MLC2v (red) and MLC2a (green) in the iPS-derived human *de novo* myocardium showed small numbers of MLC2a^pos^ cells in clusters (left) or as single cells (right). **E)** Quantification of MLC2v^pos^ CMs (red) and MLC2a^pos^ CMs (green) given as % positive CM area in the grafts after 2 weeks (2w) and 3 months (3m), suggests progressing CM maturation. Mean ± SEM. **F)** Anti CX43 staining suggests functional coupling of host myocardium with donor cardiomyocytes; human-specific cTNI (yellow), SA (red), CX43 (green). **G)** The iPSC-derived human myocardium (human-specific cTNI, green and anti-mCherry, orange) contains capillaries formed by host-derived (anti-mCherry^neg^) endothelial cells (CD31, red, see white arrows). All nuclei were stained with DAPI (blue); Scale bars 5 mm in B; 100 µm in C-F.

### hiCMA transplantation leads to substantial, albeit hiPSC line-dependent, recovery of heart function after sub-acute myocardial infarction

Cardiac MRI was performed at four time points with a long-term follow-up for 3 months (Fig. S3A). In the experimental groups B1, B2, and C, a total of 10 animals underwent coronary artery ligation to induce MI. A subset of the animals was treated with *Amber*-hiCMA transplantation (group B1, n=3) or *Ruby*-hiCMAs (group B2, n=3), while the remaining animals served as sham control group (Fig. S3B).

An unbiased assessment of the combined treatment groups B1+B2 (n=6) compared to the sham control group C (n= 4) is shown in Fig. S4A with a trend for an increased LV-rEF after 3 months. Including only noticeably infarcted animals in figure 5A, the values for the relative LV-rEF confirmed a significant decrease in contractility after myocardial infarction for the cell-treated group B (0.78 ± 0.08 vs. pre-MI (= 1); p<0.05 for n=5), and for the untreated group C animals (0.70 ± 0.10 vs. pre-MI (= 1); p<0.05, for n=3). Already on day 14 increased recovery of heart function compared to the control group became apparent for the cell-treated group B animals (LV-rEF of 0.83 ± 0.11). More pronounced improvement of heart function was observed after 3 months (LV-rEF of 0.85 ± 0.03), which was significantly higher than for the sham control animals (0.73 ± 0.03; adjusted p value < 0.05 for unpaired t test with Holm-Ṧidák method).

**Figure 5:**
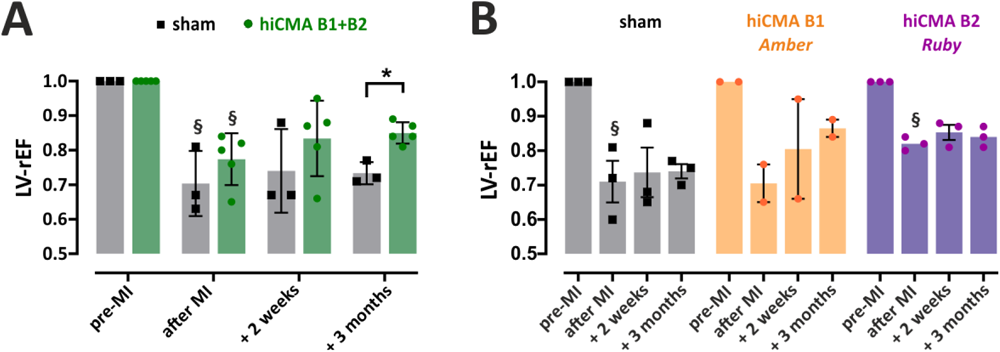
hiCMA transplantation can lead to substantial recovery of heart function. **A)** LV-EF values relative to pre-MI values (LV-rEF) were used to assess the outcome of individual hiCMA-treated animals. Only animals with a significant reduction of LV-rEF two weeks after coronary ligation were included (§, p < 0.05 for paired t-test vs pre-MI) i.e. the treatment groups B1+B2 (green symbols) with n = 5 (#06, #07, #10, #12, #13) and the sham control group C (black symbols) with n = 3 (#09, #11, #15). This direct comparison showed a significantly increased LV-rEF of 0.85 ± 0.03 vs. 0.73 ± 0.03 for treated vs. sham-treated animals, respectively, after 3 months (*, adjusted p value < 0.05 for unpaired t test with Holm-Ṧidák method and n = 3-5). **B)** This data was re-analyzed for treatment groups B1 (#06, #07: orange symbols) and B2 (#10, #12, #13; purple symbols), separately. While the low number of treated animals did not allow achieving significance, a clear trend towards increased relative ejection fraction two weeks and 3 months after cell injection was observed after hiCMA treatment using hiCMA from MHHi0001-A-11 ‘Amber’ indicating substantial recovery of myocardial contractility after treatment. In contrast, no improvement was observed for hiCMA treatment using iPSC line MHHi0001-A-5 ‘Ruby’.

Finally, a cell line dependent analysis revealed that only animals in group B1, which received *Amber*-hiCMAs, contributed to the increased recovery of heart function compared to control animals (LV-rEF 0.81 ± 0.21 vs. 0.71 ± 0.01 at baseline 14d after MI; LV-rEF 0.87 ± 0.04 three months after MI) (Fig. 5B). In contrast, animals that received *Ruby*-hiCMAs obviously did not contribute to the significantly increased functional improvement in group B animals after three months, potentially reflecting the poor *in vitro* contractility of the respective *Amber*-and *Ruby*-CM lots assessed by the BCT assay (Fig. S1H). In individual animals, the extent of functional improvement (LV-rEF change after 3 months) seemed to correlate with the extent of initial loss of contractility 2 weeks after MI (Fig. S4B).

The presence of non-contractile scar tissue demonstrated by HE staining (Fig. S4C) was confirmed by cardiac MRI with characteristic values of T1-relaxation times and an increased extracellular volume (ECV) of 80% showing fibrotic tissue changes in the scar (vs. an ECV of ∼25% in the remote region (Fig. S4D). Scar tissue characteristics did not differ between the treated and untreated group, however, the size of the fibrotic area was not determined.

### hiCMA transplantation is associated with a transient increase in ventricular arrhythmia

Telemetric ECG recording was performed to assess the occurrence of described arrhythmias ^10, 11^. All animals received an initial anti-arrhythmic medication in the peri-infarct period. For each arrhythmia type (Fig. 6A), the burden was quantified as number of events per day (Fig. 6B,C). During the first two weeks after MI, mainly premature ventricular complexes (PVCs), ventricular couplets, ventricular triplets, and – to a lesser extent – non-sustained AIVR and non-sustained VT were seen; no relevant burden of sustained AIVR or sustained VT was observed (Fig. 6B,C). hiCMA application two weeks post MI induced a distinct increase of mainly PVCs, ventricular couplets, ventricular triplets and sustained/ non-sustained AIVR. Whereas animal #06 showed an increase of arrhythmias as early as two days after hiCMA injection from *Amber*-iPSCs, animals #05, #07 (treated with the same cell line) and #10 (treated with *Ruby*-hiCMAs) exhibited no significant arrhythmogenic effect within the first 10 days after treatment. A relevant burden of non-sustained VT and some instances of sustained VT were observed in only one animal (#05) three weeks after cell injection (Fig. 6B,D).

**Figure 6:**
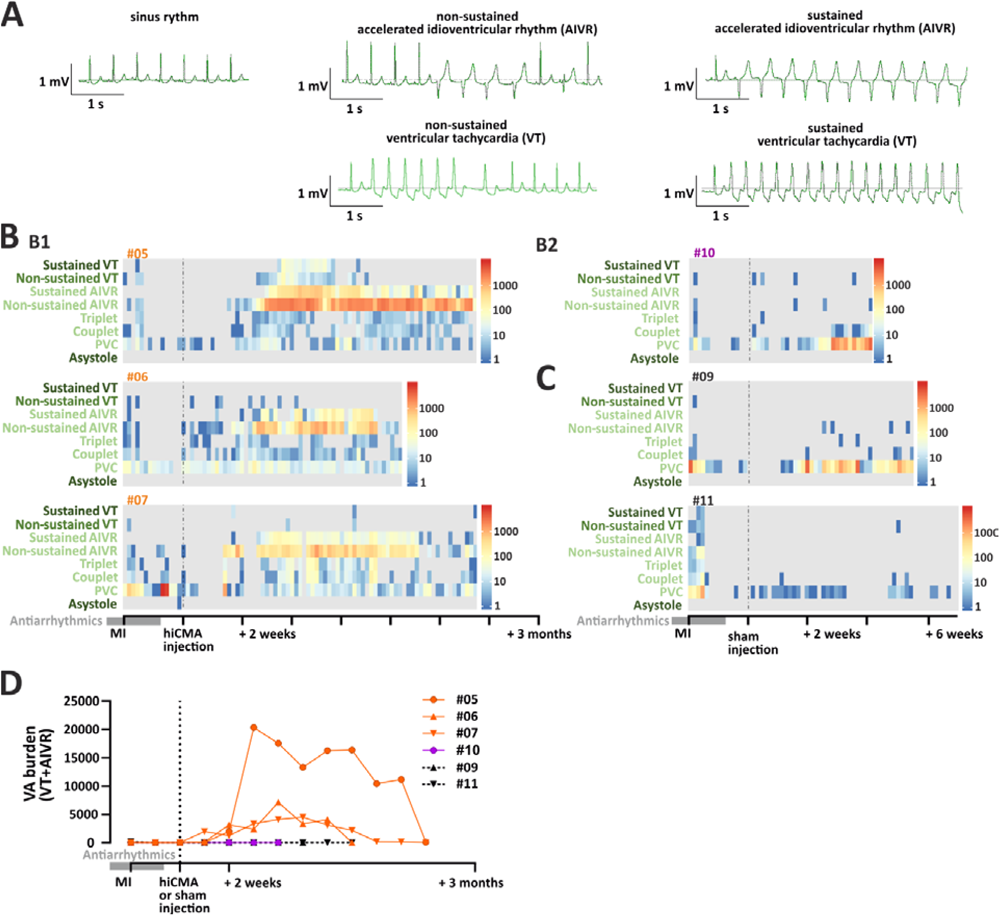
hiCMA transplantation is associated with transient graft-induced arrhythmias that substantially decline over time. **A)** Telemetric ECG recording allowed for the detection of various types of arrhythmias, including non-sustained and sustained accelerated idioventricular rhythm (AIVR), non-sustained and sustained ventricular tachycardia (VT) (see original traces). **B)** Quantitative analyses of VTs, AIVRs, triplets, couplets, premature ventricular contractions (PVCs) and events of asystole revealed the number of events per day for each type of arrhythmia and is shown as heat maps for individual animals in group B1 (#05, #06, #07) and B2 (#10) after hiCMA transplantation over time. Notably, the most severe types of arrhythmia (indicated by darker fonts’ color) were the least frequent ones. **C)** Control animals from the sham-treated group C (#09, #11) showed a significantly lower number of events per day for each type of arrhythmia. **D)** Summarized data for the ventricular arrhythmia burden (VA, i.e. VT and AIVR) per week for individual animals after hiCMA transplantation (#05, #06, #07 in orange; #10 in purple) compared to untreated control animals (#09, #11 in black).

In all treated animals, the burden of ventricular arrhythmias (AIVR+VT) peaked at 4-6 weeks after hiCMA transplantation, followed by steady decrease (Figure 6D). In contrast, we did not observe a relevant burden of arrhythmia for the PBS treated control animals #09 and #11 (Fig. 6C,D) and no relevant bradyarrhythmias (including asystoles) were detected in all respective animals.

## Discussion

For patients suffering from ischemia/reperfusion-induced heart failure, cell therapies based on iPSCs aiming at the *de novo* generation of heart muscle open new perspectives for heart repair ^22, 23^. This study addressed a number of challenges and open questions on the road to clinical translation, including the need for i) reliable cell production, ii) suitable pre-clinical animal models and ways to deal with their limitations, and iii) solutions for the safe and efficient cell application. Building on our previously developed suspension culture technologies ^3, 5^, we have produced the clinically relevant numbers of hiPSC CMs at chemically defined and in principle GMP-compliant conditions. The hiPSC line independent CMs purity of typically about 95-98% achieved by our protocol meets the requirement of at least 90% CMs content in the envisioned cell therapy product, as we have discussed in informal meetings with the competent regulatory authorities. Besides the CMs purity aspect, complementary *in vitro* assays showing the lack of persistent pluripotent cells and respective toxicological in vivo studies complementing our functional assessment in NHPs are required, which are underway. This progress paves the way towards first in man (FIM) studies, the ultimate aim of this translational research.

The generated cardiomyocyte aggregates were relatively homogeneous regarding their size patterns and CMs content across numerous independent production lots, and remained phenotypical stable for up to 5 weeks of culture maintenance, rendering cryopreservation unnecessary. This enables the spatial and temporal uncoupling of hiCMA production from their transplantation, supporting the logistics of future cardiac cell therapies. However, comparing the individual CM batches we noticed that, despite very similar differentiation efficiencies and expression patterns of the two reporter lines, our BCT *in vitro* assay showed significantly different, cell line-dependent force levels. This seems surprising as both lines were derived from the same parental cell line MHHi0001-A, and modified with the same procedures to express Venus or RedStar+GCaMP6f (in *Amber* or *Ruby* cells, respectively). For unrelated iPSC lines, an intra-batch variability of up to 20% and an inter-batch variability of ∼30% for forces of iPSC-CM derived engineered tissue was reported ^24^. Calcium sensors such as GCaMP can act as a calcium buffer ^25^, potentially leading to reduced contraction forces, however, Saleem et al. did not observe this phenomenon in engineered heart tissue ^26^. Our *in vitro* observations raised the question whether the cell line-specific hiCMA batches would also perform differently *in vivo* after transplantation.

Our NHP model of sub-acute MI was developed to explore the potential of hiCMA injection for heart function recovery in cynomolgus monkeys. In these animals, collateral myocardial blood flow is low and does not significantly influence infarct size ^27^ and potentially also recovery. Similar to Shiba et al. ^6^, Chong et al. ^10^, and Liu et al. ^11^ we have decided to inject hiCMAs at sub-acute conditions, i.e. only two weeks after experimental MI induction. This reduces the risk of cell depletion due to acute inflammation directly post MI and promotes cells engraftment prior to scar formation. From a clinical perspective, a two weeks’ timeframe is certainly not compatible with the patient-specific (autologous) production process of CMs, including differentiation, quality controls, etc., even if a stock of cryopreserved patient-specific iPSCs would be available. Therefore, the concept of cellular therapy after sub-acute MI is rather compatible with an allogeneic transplantation. This may potentially imply HLA-matched-, HLA-homozygote-or genetically engineered cells with decreased immunogenicity, but probably still requiring some degree of immunosuppression.

Our data confirm that proper immunosuppression enables the long term survival and structural integration of human CMs in macaques for at least 3 months, as reported ^10, 11^. We demonstrate that human CMs can be injected in the form of distinct cell aggregates, which are remodeled *in vivo* to form larger grafts, potentially via matrix remodeling, cell migration, cell coupling and potentially also cell proliferation.

The appearance of host-derived endothelial cells in the human grafts already after 2 weeks suggested fast vascularization of the aggregate-based grafts. Notably, the largest and most mature grafts were observed in areas of healthy donor myocardium and cardiac MRI demonstrated the presence of residual scar tissue ^28, 29^ after cell treatment. This could be a result of impaired survival in scar tissue, therefore an optimized application and targeting to the infarct border zones and the scar area might need to be explored to achieve an improved remuscularization of the injured myocardium.

Published data suggested the substantial improvement of heart function and substantial hPSC-derived donor grafts after transplantation of relatively large numbers of ∼10^9^ single cell hESC-derived CMs per NHP heart ^11^. At the same time, previous studies by us and others, suggested a substantial cell loss immediately after cell injection by backflush from the injection channel and in particular the venous route ^30, 31^. Therefore, we hypothesized that the intra-myocardial injection of cell aggregates, rather than single cells, may support cell retention and engraftment efficiency. Indeed, our application of CM aggregates of ∼200 µm diameter led to the formation of relatively large grafts despite using relatively low cell numbers of 5 x10^7^ CMs / NHP heart. However, due to the limited animal numbers that can be tested due to ethical, regulatory and financial restrictions, we have not performed a direct comparison of our aggregate approach versus single cell application.

The small numbers of animals usually included in NHP studies led to additional challenges for functional analyses; already pre-MI LVEF values showed a substantial variation. Part of the variance is due to inter-individual differences in age, weight and coronary artery anatomy, in addition, two animals were evaluated using the Real-Time approach due to distorted ECG signals, while standard CINE MRI was used for all other animals ^18^. Consequently, we decided to determine heart function as LV-rEF, i.e. relative to the pre-MI LV-EF, as this allowed for comparative analyses in small groups of large animals. Experimental MI was successful in most animals with a loss of contractile myocardium in left ventricular and septal areas near the apex, the extent of LV-rEF reduction varied based on the animals’ anatomy and positioning of the coronary artery ligation. We found that in our animal model, serum levels of TnT can be used as a valuable biomarker not only for the presence of myocardial damage, as described by Fredericks et al. ^32^, but also for loss of contractile function. Total CK and CK-MB were poorer markers, most likely because of the reduced cardiac specificity of CK-MB in cynomolgus monkeys compared to humans ^32^.

While the generated MRI data suggest improvement of heart function compared to sham treated animals already on d14 after treatment, significant improvement of heart function was observed for the cell-treated group B animals after 3 months. Obviously, however, only animals that received hiCMAs produced from the *Amber* reporter line contributed to this significant functional improvement; no increase in LVEF was observed in animals that received hiCMAs from the *Ruby* reporter line. Since *Amber*-BCTs outperformed *Ruby*-BCTs with respect to their contractile force development *in vitro*, it is tempting to speculate that the *Amber*-hiCMAs, when engrafted in the post-infarcted NHP heart, promoted heart function recovery by their contractile properties as mechanism of action, which is in line with recent data by Stüdemann et al. who provided a clear link between the transplanted CM’s contractility and the recipient’s heart function in a guinea pig model ^33^. However, we cannot exclude a contribution of paracrine or other indirect mechanisms discussed in the field ^34–36^. Clearly, this hypothesis needs to be fortified by additional future studies. Nonetheless, our observations clearly highlight both the requirement for a suitable *in vitro* potency assay as requested by regulatory authorities ^37, 38^, and the utility of our BCT approach as a practical basis for potency testing.

Clearly, besides the efficacy of a stem cell-based treatment, the patients’ safety will be of utmost importance for clinical translation. In this NHP study we confirm the occurrence of ventricular arrhythmias as described in the literature ^10, 11^, which were transient and non-life-threatening. Similar to Liu et al. ^11^, we did not observe a negative effect by the reporter gene GCaMP and apparently the application of CMs in the form of aggregates instead of single cells posed no increased risk of arrhythmias. At this point, we can only speculate that the injection of hiCMAs and their efficient engraftment should lead to less off-target effects because fewer single cells are lost to circulation. Following promising results in guinea pig and pigs ^39^, recently a first clinical trial for hiPSC-CM-based engineered cardiac patches was initiated in Germany/ Europe, (BioVAT-HF, ClinicalTrials.gov Identifier: NCT04396899). While this delivery approach is not suffering from limited cell retention, it is associated with other challenges, such as the limitation of superficial tissue placement. It will be exciting to see how the different treatment strategies will perform in terms of ATMP testing, i.e. safety and also efficacy of clinical application.

## Conclusion

We show that the transplantation of hiCMAs resulted in the formation of substantial, structurally integrated muscular grafts and improved heart function in an NHP model of sub-acute MI. The application of CM aggregates, rather that single cells, enabled the usage of relatively small cell numbers, thereby promoting the efficiency of the procedure by reducing cell production costs and clinical risks associated with high cell number administration and accompanying cell death. According to this study, hiCMA injection can be considered reasonably safe, however, for clinical translation efficient strategies for the pharmacological prevention and treatment of arrhythmias are of utmost importance; in addition, temporary implantation of a cardioverter-defibrillator (ICD) could be considered. Since we also demonstrate the temporal uncoupling of cell production from transplantation and the implementation of an *in vitro* potency test, our PoC study provides pragmatic new avenues for hiPSC-based heart repair.

## Acknowledgements

We would like to thank Kevin Ullmann experimental support in cardiomyocytes production, Doreen Lenz and Ingrid Schmidt-Richter for excellent technical assistance, Tamara Becker, Sophie Mißbach for support with animal handling, Christian Hackenbroich and Ulrich Molitoris for support with animal anesthesia, Hasti Moussavi for support with MRI analysis, Martina Bleyer and Evelyn Gruber-Dujardin for support with animal pathology, Cristina Luminita Baciu and Daniel Reckel for their help with data handling and analysis.

## Supplemental Material

Supplemental Methods and Supplemental Figures S1-S4 Supplemental Videos S1 & S2

## References

1. McDonagh TA, Metra M, Adamo M, Gardner RS, Baumbach A, Böhm M, et al. 2021 ESC Guidelines for the diagnosis and treatment of acute and chronic heart failure: Developed by the Task Force for the diagnosis and treatment of acute and chronic heart failure of the European Society of Cardiology (ESC). With the special contribution of the Heart Failure Association (HFA) of the ESC. Eur J Heart Fail 2022;24(1):4-131.

2. Martin U. Genome stability of programmed stem cell products. Adv Drug Deliv Rev 2017;120:108–117.

3. Manstein F, Ullmann K, Kropp C, Halloin C, Triebert W, Franke A, et al. High density bioprocessing of human pluripotent stem cells by metabolic control and in silico modeling. Stem Cells Transl Med 2021.

4. Kempf H, Kropp C, Olmer R, Martin U, Zweigerdt R. Cardiac differentiation of human pluripotent stem cells in scalable suspension culture. Nat Protoc 2015;10(9):1345–61.

5. Halloin C, Schwanke K, Lobel W, Franke A, Szepes M, Biswanath S, et al. Continuous WNT Control Enables Advanced hPSC Cardiac Processing and Prognostic Surface Marker Identification in Chemically Defined Suspension Culture. Stem Cell Reports 2019;13(2):366–379.

6. Shiba Y, Fernandes S, Zhu WZ, Filice D, Muskheli V, Kim J, et al. Human ES-cell-derived cardiomyocytes electrically couple and suppress arrhythmias in injured hearts. Nature 2012;489(7415):322-5.

7. Laflamme MA, Chen KY, Naumova AV, Muskheli V, Fugate JA, Dupras SK, et al. Cardiomyocytes derived from human embryonic stem cells in pro-survival factors enhance function of infarcted rat hearts. Nat Biotechnol 2007;25(9):1015-24.

8. van Laake LW, Passier R, Monshouwer-Kloots J, Verkleij AJ, Lips DJ, Freund C, et al. Human embryonic stem cell-derived cardiomyocytes survive and mature in the mouse heart and transiently improve function after myocardial infarction. Stem Cell Res 2007;1(1):9–24.

9. Romagnuolo R, Masoudpour H, Porta-Sánchez A, Qiang B, Barry J, Laskary A, et al. Human Embryonic Stem Cell-Derived Cardiomyocytes Regenerate the Infarcted Pig Heart but Induce Ventricular Tachyarrhythmias. Stem Cell Reports 2019;12(5):967–981.

10. Chong JJ, Yang X, Don CW, Minami E, Liu YW, Weyers JJ, et al. Human embryonic-stem-cell-derived cardiomyocytes regenerate non-human primate hearts. Nature 2014;510(7504):273-7.

11. Liu YW, Chen B, Yang X, Fugate JA, Kalucki FA, Futakuchi-Tsuchida A, et al. Human embryonic stem cell-derived cardiomyocytes restore function in infarcted hearts of non-human primates. Nat Biotechnol 2018;36(7):597–605.

12. Shiba Y, Gomibuchi T, Seto T, Wada Y, Ichimura H, Tanaka Y, et al. Allogeneic transplantation of iPS cell-derived cardiomyocytes regenerates primate hearts. Nature 2016;538(7625):388-391.

13. Amato R, Gardin JF, Tooze JA, Cline JM. Organ Weights in Relation to Age and Sex in Cynomolgus Monkeys (Macaca fascicularis). Toxicol Pathol 2022;50(5):574–590.

14. Nagai T, Ibata K, Park ES, Kubota M, Mikoshiba K, Miyawaki A. A variant of yellow fluorescent protein with fast and efficient maturation for cell-biological applications. Nat Biotechnol 2002;20(1):87–90.

15. Haase A, Kohrn T, Fricke V, Ricci Signorini ME, Witte M, Göhring G, et al. Establishment of MHHi001-A-5, a GCaMP6f and RedStar dual reporter human iPSC line for in vitro and in vivo characterization and in situ tracing of iPSC derivatives. Stem Cell Res 2021;52:102206.

16. Kensah G, Roa Lara A, Dahlmann J, Zweigerdt R, Schwanke K, Hegermann J, et al. Murine and human pluripotent stem cell-derived cardiac bodies form contractile myocardial tissue in vitro. Eur Heart J 2013;34(15):1134–46.

17. Haggerty HG, Proctor SJ. Chronic administration of belatacept, a T-cell costimulatory signal blocker, in cynomolgus monkeys. Toxicol Sci 2012;127(1):159–68.

18. Moussavi A, Mißbach S, Serrano Ferrel C, Ghasemipour H, Kötz K, Drummer C, et al. Comparison of cine and real-time cardiac MRI in rhesus macaques. Sci Rep 2021;11(1):10713.

19. Wang X, Joseph AA, Kalentev O, Merboldt KD, Voit D, Roeloffs VB, et al. High-resolution myocardial T(1) mapping using single-shot inversion recovery fast low-angle shot MRI with radial undersampling and iterative reconstruction. Br J Radiol 2016;89(1068):20160255.

20. Weir JP. Quantifying test-retest reliability using the intraclass correlation coefficient and the SEM. J Strength Cond Res 2005;19(1):231–40.

21. Martin Bland J, Altman D. Statistical methods for assessing agreement between two methods of clinical measurement. The Lancet 1986;327(8476):307-310.

22. Braunwald E. Cardiac cell therapy: a call for action. Eur Heart J 2022;43(25):2352–2353.

23. Menasché P. Cell therapy trials for heart regeneration -lessons learned and future directions. Nat Rev Cardiol 2018;15(11):659–671.

24. Mannhardt I, Saleem U, Mosqueira D, Loos MF, Ulmer BM, Lemoine MD, et al. Comparison of 10 Control hPSC Lines for Drug Screening in an Engineered Heart Tissue Format. Stem Cell Reports 2020;15(4):983–998.

25. McMahon SM, Jackson MB. An Inconvenient Truth: Calcium Sensors Are Calcium Buffers. Trends Neurosci 2018;41(12):880–884.

26. Saleem U, Mannhardt I, Braren I, Denning C, Eschenhagen T, Hansen A. Force and Calcium Transients Analysis in Human Engineered Heart Tissues Reveals Positive Force-Frequency Relation at Physiological Frequency. Stem Cell Reports 2020;14(2):312–324.

27. Yang XM, Liu Y, Liu Y, Tandon N, Kambayashi J, Downey JM, et al. Attenuation of infarction in cynomolgus monkeys: preconditioning and postconditioning. Basic Res Cardiol 2010;105(1):119–28.

28. Stuckey DJ, McSweeney SJ, Thin MZ, Habib J, Price AN, Fiedler LR, et al. T₁ mapping detects pharmacological retardation of diffuse cardiac fibrosis in mouse pressure-overload hypertrophy. Circ Cardiovasc Imaging 2014;7(2):240–9.

29. Haaf P, Garg P, Messroghli DR, Broadbent DA, Greenwood JP, Plein S. Cardiac T1 Mapping and Extracellular Volume (ECV) in clinical practice: a comprehensive review. J Cardiovasc Magn Reson 2016;18(1):89.

30. Martens A, Rojas SV, Baraki H, Rathert C, Schecker N, Zweigerdt R, et al. Substantial early loss of induced pluripotent stem cells following transplantation in myocardial infarction. Artif Organs 2014;38(11):978–84.

31. van den Akker F, Feyen DA, van den Hoogen P, van Laake LW, van Eeuwijk EC, Hoefer I, et al. Intramyocardial stem cell injection: go(ne) with the flow. Eur Heart J 2017;38(3):184–186.

32. Fredericks S, Merton GK, Lerena MJ, Heining P, Carter ND, Holt DW. Cardiac troponins and creatine kinase content of striated muscle in common laboratory animals. Clin Chim Acta 2001;304(1-2):65–74.

33. Stüdemann T, Rössinger J, Manthey C, Geertz B, Srikantharajah R, von Bibra C, et al. Contractile Force of Transplanted Cardiomyocytes Actively Supports Heart Function After Injury. Circulation 2022:101161circulationaha122060124.

34. Tachibana A, Santoso MR, Mahmoudi M, Shukla P, Wang L, Bennett M, et al. Paracrine Effects of the Pluripotent Stem Cell-Derived Cardiac Myocytes Salvage the Injured Myocardium. Circ Res 2017;121(6):e22–e36.

35. El Harane N, Kervadec A, Bellamy V, Pidial L, Neametalla HJ, Perier MC, et al. Acellular therapeutic approach for heart failure: in vitro production of extracellular vesicles from human cardiovascular progenitors. Eur Heart J 2018;39(20):1835–1847.

36. Zhu K, Wu Q, Ni C, Zhang P, Zhong Z, Wu Y, et al. Lack of Remuscularization Following Transplantation of Human Embryonic Stem Cell-Derived Cardiovascular Progenitor Cells in Infarcted Nonhuman Primates. Circ Res 2018;122(7):958–969.

37. Bravery CA, Carmen J, Fong T, Oprea W, Hoogendoorn KH, Woda J, et al. Potency assay development for cellular therapy products: an ISCT review of the requirements and experiences in the industry. Cytotherapy 2013;15(1):9–19.

38. Tsokas K, McFarland R, Burke C, Lynch JL, Bollenbach T, Callaway DA, 2nd, et al. Reducing Risks and Delays in the Translation of Cell and Gene Therapy Innovations into Regulated Products. NAM Perspect 2019;2019.

39. Querdel E, Reinsch M, Castro L, Köse D, Bähr A, Reich S, et al. Human Engineered Heart Tissue Patches Remuscularize the Injured Heart in a Dose-Dependent Manner. Circulation 2021;143(20):1991–2006.

