## Supplemental Material for "Cell therapy with human iPSC-derived cardiomyocyte aggregates leads to efficient engraftment and functional recovery after myocardial infarction in non-human primates"

*[8] Fraunhofer Institute for Toxicology and Experimental Medicine, Hannover Medical School, Germany*

### Supplemental methods

#### hiPSC culture, differentiation and analysis of cardiomyocyte aggregates

Human iPSC cultivation, chemically defined cardiac differentiation, and cell/ aggregate analysis was performed essentially as described [1, 2]. Monolayer cultures of the hiPSCs reporter lines MHHi0001-A-11 (‘*Amber*’, constitutively expressing Venus(nucmem) [3] under control of the CAG promoter from the AAVS1 locus) or MHHi0001-A-5 (‘*Ruby*’, constitutively expressing RedStar(nucmem) and the Calcium sensor GCaMP6f under control of the CAG promoter from the AAVS1 locus [4] were expanded in E8 medium on Geltrex-coated T75 and T175 flasks. For single cell inoculation in suspension culture, cells were dissociated by 3-5 min of Accutase (Thermo Fisher Scientific) treatment and seeded at 0.15 x $10^6$cells/ml in 300 ml of E8 medium supplemented with 10 mM Y-27632 (Rock-Inhibitor, RI; [5] and 0.1% Pluronic-F68 solution (Gibco, PLU) into 500 mL Disposable Spinner Flask with Vent Cap (Corning), stirred at 60 RPM for 48h.

Differentiation of the resulting hiPSC aggregates was initiated in CDM3 medium supplemented with 5 µM CHIR99021 (Tocris Bioscience) in the same Spinner platform. Precisely 24h later, the medium was replaced by fresh CDM3 supplemented with 5 µM IWP2 (Tocris Bioscience). Precisely 48h later, CDM3-IWP2 was replaced with pure CDM3 and 75% medium was replaced every 2–3 days thereafter. After check point 1 (CP1; see Fig. 1) of differentiation, resulting cardiomyocyte aggregates (hiCMAs) were cultured in $RB^+$ medium (RPMI-B27 with insulin, Gibco with Penicillin/Streptomycin, Sigma-Aldrich) replaced every 3-4 days until check point 2 (CP2; see Fig. 1).

To monitor the hiCMA diameters, up to twelve independent light microscopic images of hiCMA samples (placed in 6-well plate) were captured at 4x magnification (Olympus CKx41 inverted microscope, Olympus) followed by automated diameter analysis via ImageJ (https://imagej.nih.gov/ij/); mean diameters represent the arithmetic average of at least 70 individual aggregates.

#### Flow cytometry

For CM content monitoring on CP1/ CP2, hiCMA samples were dissociated by STEMdiff Cardiomyocyte Dissociation Kit (Stem Cell Technologies) following suppliers instructions. For flow cytometry (FC) analysis, cells were treated with FIX&PERM Solution (Thermo Fisher Scientific) and stained with respective antibodies specific to anti-cardiac Troponin T (cTnT; 1:200, clone 13-11, Thermo Fisher Scientific), anti-sarcomeric α-Actinin (1:800, EA53, Sigma-Aldrich or 1:20, REA402, Miltenyi Biotech), anti-Myosin Heavy Chain (1:20, MF20, DSHB), or isotype control mouse-IgG1 (1:200, DAKO). The primary antibodies were detected by appropriate Cy5-conjugated antibodies (1:200, Jackson ImmunoResearch) followed by data acquisition on a MACSquant Analyzer 10 Flow Cytometer (Miltenyi Biotec) and data analysis by FlowJo software (Flowjo, LLC).

**Generation of bioartificial cardiac tissue and force measurement**

For the generation of BCTs, the hiCMAs were dissociated and 1 million CMs were used for tissue preparation as described [6]. CMs were mixed with 10% human dermal fibroblasts (ATCC) in 100 µl BCT medium [6] per tissue. An extracellular matrix mixture (i.e. 150 µl/ BCT) composed of 1.35 mg/ml rat collagen type I (Trevigen), 10% Geltrex (Life Technologies), and 3.8% 0.4 M NaOH was added. The cell-matrix mixture was poured into a custom-made silicon mold containing two titanium rods (distance 6 mm; initial slack length) and solidified at 37 °C for 30 minutes. Then, the construct was covered by 5 ml BCT medium with 30 µM L-ascorbic acid (Sigma-Aldrich) and cultured under standard cell culture conditions with medium change every 1 or 2 days. For all BCTs, growing static stretch (G-stretch) was applied through stepwise elongation by 400 µm on days 7 and 11. Tissue morphology was monitored during cultivation using an AxioObserver Z1 fluorescence microscope and ZEN software from Zeiss [4]. Mechanical forces of the tissues were measured on day 14 of tissue cultivation (equivalent to CP3 *in vivo*) in BCT medium at 37 °C, 5% CO2 levels using a custom-made bioreactor system and analysis software (Central Research Workshop, Hannover Medical School) as previously described [22]. Active contraction force was determined at increasing preload (in 100 or 200 µm increments until 1 mm in total) by measuring the response to electric stimulation (5x) with biphasic pulses (10 ms, ±25 V). Passive force for each preload step was defined by the difference between baselines at the actual step and at the original length (L0).

For video-optical calcium analysis, videos at a sampling rate of 47 frames/s for 15 sec were acquired using the AxioObserver Z1 fluorescence microscope (Zeiss, Jena, DE). Changes in fluorescence intensity while spontaneous contractions were calculated using the ZEN module Physiology (Dynamics) of the ZENpro software (Zeiss, Jena, DE). After selection of a Region of Interest (ROI) and defining an area for background correction, the ratio calculation for the fluorescence channel was performed as previously described by Tian, Martinez et al. [7].

**Cell preparation for transplantation**

For transplantation, hiCMA were collected from the culture medium by 1 min centrifugation at 300g. After media removal, the hiCMA pellet was carefully washed with DPBS (Gibco), re-collected by centrifugation and carefully filled-up with DPBS to a total volume of 800 µl in scaled 1.5 ml tubes (StarLab). Each hiCMA sample injected per heart contained ~50 million CMs (calculated based on the analysis of parallel, representative hiCMA aliquots dissociated into single cells for counting). For injection, the ~ 800 µl of hiCMAs/ DPBS suspension was carefully transferred to a 26G syringe.

**Animal anesthesia**

For all major surgery, animals were pre-medicated with glycopyrronium bromide, medetomidine, maropitant, and meloxicam, then anaesthetized with ketamine (10 to 15 mg $kg^{-1}$) and propofol, (0.4 to 0.6 mg $kg^{-1}$ $min^{-1}$), intubated and ventilated using isoflurane to maintain anesthesia. Perioperative and postoperative analgesia was performed with fentanyl (0.1 to 0.5 µg $kg^{-1}$ $min^{-1}$ i.v.), buprenorphine (5-10 µg $kg^{-1}$ s.c.) and bupivacaine hydrochloride (1 mg $kg^{-1}$, intercostal nerve blockade).

#### Echocardiography

Trans-esophageal cardiac echocardiography was performed before and after induction of infarction under general anesthesia. Images were captured with an EPIQ 7G device using an S7-3t trans-esophageal phased ultrasound transducer as four–chamber, two-chamber and short axis views. Functional analysis was performed using a CX50 device and software version OS-12.676 (all from Philips N.V. Amsterdam, Netherlands).

#### ECG monitoring

Continuous monitoring for clinically relevant arrhythmias was established using an implantable telemetry device (PhysioTel Digital M01, Data Sciences International (DSI), St. Paul, MN, USA). The device was implanted subcutaneously in the abdominal region at the time of LAD ligation surgery during general anesthesia. Leads were tunneled to achieve a lead II configuration and a single lead ECG was continuously recorded from day 0 (induction of myocardial infarction) until sacrification of the animal. ECG recordings were analyzed with focus on relevant bradyarrhythmias and tachyarrhythmias and automated results were reviewed by a physician. DSI Ponemah® version 6.20 with Pattern Recognition Option (DSI, St. Paul, MN, USA) was used for ECG telemetry data collection and analysis. Ventricular arrhythmias were classified as premature ventricular complexes (PVCs; QRS duration >60 ms), ventricular couplets, ventricular triplets, non-sustained accelerated idioventricular rhythms (AIVR; heart rate <180 bpm), sustained AIVR, non-sustained ventricular tachycardia (VT; heart rate ≥180 bpm) or sustained VT as described by Liu et al. [8]. AIVR and VT were considered sustained if ≥30 conjugated respective wide complexes occurred. Representative ECG tracings are shown in Figure 6A. Arrhythmia burden was quantified as number of events per day for each arrhythmia type as well as for the total ventricular arrhythmia burden (aggregate of VT and AIVR).

#### Cardiac MRI

The monkeys underwent MRI measurements before myocardial infraction (pre-MI), before cell transplantation (baseline), and 2 and 12 weeks after cell transplantation (Fig. 3A). All experiments were performed on a 3T MR-system (Magnetom Prisma, Siemens Healthineers, Erlangen, Germany) using a 16-channel multipurpose coil (Variety, Noras MRI products,

Hoechberg, Germany) for signal reception. The anesthetized and intubated monkeys were mechanically ventilated (Servo Ventilator 900C, Siemens-Elema AB, Sweden) and physiological parameters (e.g., heart rate, breathing rate, and end-tidal $CO_2$) were continuously monitored.

For the assessment of the left ventricular function, short-axis images were acquired during forced expiratory breath holds using an ECG-gated segmented cine Fast Low-Angle Shot (FLASH) technique (repetition time (TR) = 6.5 ms, echo time (TE) = 2.8 ms, flip angle = 12°, bandwidth = 215 Hz/Px, field of view (FOV) = 132 × 149 mm², 6 segments, spatial resolution = 0.85 × 0.85 × 3 $mm^3$, 13–17 slices covering the heart from the apex to the base with an acquisition time of 15-20 seconds per slice) as previously described [9]. In addition, Real-Time MRI during free breathing was acquired using spoiled radial FLASH (TR = 2.9 ms, TE = 1.9 ms, flip angle = 8°, bandwidth = 1305 Hz/Px, FOV = 128 × 128 mm², base resolution = 128, 13 spokes per frame, Nyquist undersampling factor = 15.5, spatial resolution = 0.9 × 0.9 × 3 $mm^3$, acquisition time per frame = 37 ms, 13-17 slices and 210 frames per slice). In case of a corrupted ECG signal (animal #05, and #10), the ventricular function was assessed using Real-Time MRI instead of Cine MRI at all four time points of these monkeys.

To assess the volume and function of the left ventricle, the myocardium was manually segmented in a blind fashion by two observers with at least two years of experience in cardiovascular MRI analyses of primates using the software package Segment (Version 2.0 R6435, Medviso, Lund, Sweden) [10]. Besides the standard volumetric and functional parameter of the left ventricle (end-diastolic volume (EDV), end-systolic volume (ESV), stroke volume (SV), ejection fraction (EF)) the relative EF (rEF) was defined as the EF normalized by the respective pre-MI EF of the same monkey. Noteworthy to mention, despite systematic differences in the estimated absolute values of EF acquired by cine and Real-Time MRI, no significant differences in rEF are expected [8].

Maps of the T1-relaxation time were acquired in the 4-chamber view before and 15 minutes after contrast agent administration (Gadobutrol (GadovistTM), 0.1 mmol/kg per BW) using a single-slice fast Real-Time single-shot inversion recovery FLASH technique (TR = 2.7 ms, TE 1.8 = ms, flip angle = 6°, bandwidth = 1345 Hz/Px, FOV = 128 × 128 mm², base resolution = 128, 15 spokes per frame, Nyquist undersampling factor = 13.4, spatial resolution = 1 × 1 × 4 $mm^3$, acquisition time per frame = 40 ms, total acquisition time = 3 s) [11]. T1-relaxation times and extracellular volumes (ECV) were estimated within the region of the scar after infarction as well as in an unaffected region of the myocardium (remote). The ECV was calculated as the hematocrit (hct) normalized ratio of pre- and post-contrast R1-relaxation constant (R1 = 1/T1) differences of the myocardium and blood, respectively (ECV = (1−hct) × (ΔR1myocardium/ΔR1blood)). A hematocrit of 0.4 was estimated for all animals.

**Histology and Immunostaining**

Adherent cells were fixed with 2% paraformaldehyde (PFA, Sigma-Aldrich) for 10 min before immunofluorescence staining. NHP hearts were processed for histology as snap frozen samples or paraffin-embedded samples. Slide-mounted hiCMAs and snap frozen heart tissue sections (both processed by embedding in Tissue-Tek O.C.T Compound (Sakura Finetek Europe) for cryo-sectioning) were fixed with 4% PFA for 4 minutes. Samples were blocked and permeabilized using Tris-buffered saline containing 0.25% Triton-X 100 and 5% donkey serum. Incubation with primary antibodies was performed either for 1 hour at RT or overnight at 4 ˚C. Secondary antibodies were added for 30 min at RT. Both primary and secondary antibodies were diluted in PBS w/o $Ca^{2+}$ and $Mg^{2+}$ + 1% bovine serum albumin (BSA) and used in ratios listed in tables S1, S2 and S3. Nuclei were stained for 15 min with 4',6-diamidino-2-phenylindole (DAPI). Images were taken using an AxioObserver A1 or Z1 fluorescence microscope and analyzed by AxioVision software 4.71 or ZEN software (both Zeiss).

**Histomorphometry**

For quantification, immunohistological images were taken with a 10x objective at the AxioObserver Z1 fluorescence microscope system and processed with ZEN 3.3 software (all Zeiss, Jena, DE). Three images per animal containing at least 1 graft area were imported into Image J software (version 1.53 c) where graft and stained areas (region of interest or ROI) were measured. Graft area was determined based on GFP or RedStar signal expressed by transgene cells that successfully engrafted in monkey hearts. Before area quantification, hyperstack images were split into single channel images with gray values and 8-bit quality. If necessary, background subtraction, brightness enhancement and median filtering was applied to images. Global threshold was adjusted using the Auto Threshold function and selecting the best fitting option (values: 9 – 199). ROI was selected and surrounding signal was deleted; the remaining area was quantified using the analyze particles function and taking the sum of all. Quantified area of MLC2a and MLC2v signal ($\mu m^2$) is represented in correlation to positive graft area (%).

**Statistical analysis**

The GraphPad Prism Version 9 (GraphPad Software, Inc., La Jolla, CA, USA) was used for statistical analyses. Unless indicated otherwise, descriptive statistical analysis was expressed as Mean ± Standard Deviation. Continuous numerical variables were compared using the student’s t-test for independent samples, significance was evaluated by two-tailed testing, and assumed at $p<0.05$.

To compare more than two groups, a two-way ANOVA with Tukey's or Bonferroni's multiple comparisons test was performed. A $p$-value $<0.05$ was considered statistically significant.

### Supplemental tables

#### Table S1: primary antibodies (Flow + IF)

| Antibody | Isotype | Company | Dilution |
|---|---|---|---|
| ACTN2 (SA) #A7811 | mouse IgG1 | Sigma-Aldrich, St. Louis, US | 1:800 |
| CD31 #M0823 | mouse IgG1 kappa | Dako, Agilent, Santa Clara, US | 1:20 |
| cTnI #ab52862 | rabbit IgG | Abcam, Cambridge, UK | 1:500 |
| CX43 #C8093 | mouse IgM | Sigma-Aldrich, St. Louis, US | 1:1000 |
| GATA4 #ab124265 | rabbit IgG | Abcam, Cambridge, UK | 1:1000 |
| GFP # R1091P | goat IgG | Acris, OriGene, Rockville, US | 1:500 |
| Ki67 #M7240 | mouse IgG1 | Dako, Agilent, Santa Clara, US | 1:100 |
| mCherry #ab167453 | rabbit IgG | Abcam, Cambridge, UK | 1:500 |
| MLC2a #311011 | mouse IgG2b | Synaptic Systems, Göttingen, DE | 1:100 |
| MLC2v #ab79935 | mouse IgG1 | Abcam, Cambridge, UK | 1:100 |
| N-Cadherin #C2542-.2ML | mouse IgG | Sigma-Aldrich, St. Louis, US | 1:100 |
| TTN (9 D10) #AB528491 | mouse IgM | Hybridoma Bank; University of Iowa, US | 1:200 / 1:20 (conc.) |
| TNNT #MS-295-P | mouse IgG1 | Thermo Fisher Scientific, Waltham, US | 1:100 |
| TNNT #15513-1-AP | rabbit IgG | Proteintech, Manchester, UK | 1:500 |

#### Table S2: Isotype control antibodies

| Antibody | Isotype | Company |
|---|---|---|
| Negative Control #X0942 | mouse IgM | Dako, Agilent, Santa Clara, US |
| Negative Control #ab37415 | rabbit IgG | Abcam, Cambridge, UK |
| Negative Control #X0943 | mouse IgG2a | Dako, Agilent, Santa Clara, US |
| Negative Control #X0944 | mouse IgG2b | Dako, Agilent, Santa Clara, US |
| Negative Control #X0931 | mouse IgG1 | Dako, Agilent, Santa Clara, US |
| Negative Control #ab37373 | Goat IgG | Abcam, Cambridge, UK |

**Table S3: Secondary antibodies**

| Host Species and Reactivity | Conjugation | Company | Dilution |
|---|---|---|---|
| Donkey anti-mouse IgG #ab175658 | Alexa Fluor 405 | Abcam, Cambridge, UK | 1:2000 |
| Donkey anti-mouse IgG #715-605-151 | Alexa Fluor 647 | Dianova, Hamburg, DE | 1:300 |
| Donkey anti-mouse IgG #715-165-150 | Cy3 | Dianova, Hamburg, DE | 1:300 |
| Donkey anti-mouse IgG #715-545-151 | Alexa Fluor 488 | Dianova, Hamburg, DE | 1:300 |
| Donkey anti-mouse IgM #715-605-140 | Alexa Fluor 647 | Dianova, Hamburg, DE | 1:300 |
| Donkey anti-mouse IgM #715-485-020 | DyLight 488 | Dianova, Hamburg, DE | 1:200 |
| Donkey anti-rabbit IgG #ab175651 | Alexa Fluor 405 | Abcam, Cambridge, UK | 1:2000 |
| Donkey anti-rabbit IgG #711-505-152 | DyLight 549 | Dianova, Hamburg, DE | 1:300 |
| Donkey anti-rabbit IgG #711-605-152 | Alexa Fluor 647 | Dianova, Hamburg, DE | 1:300 |
| Donkey anti-rabbit IgG #711-545-152 | Alexa Fluor 488 | Dianova, Hamburg, DE | 1:300 |
| Donkey anti-goat IgG #705-495-147 | DyLight 649 | Dianova, Hamburg, DE | 1:300 |

### Supplemental Figures

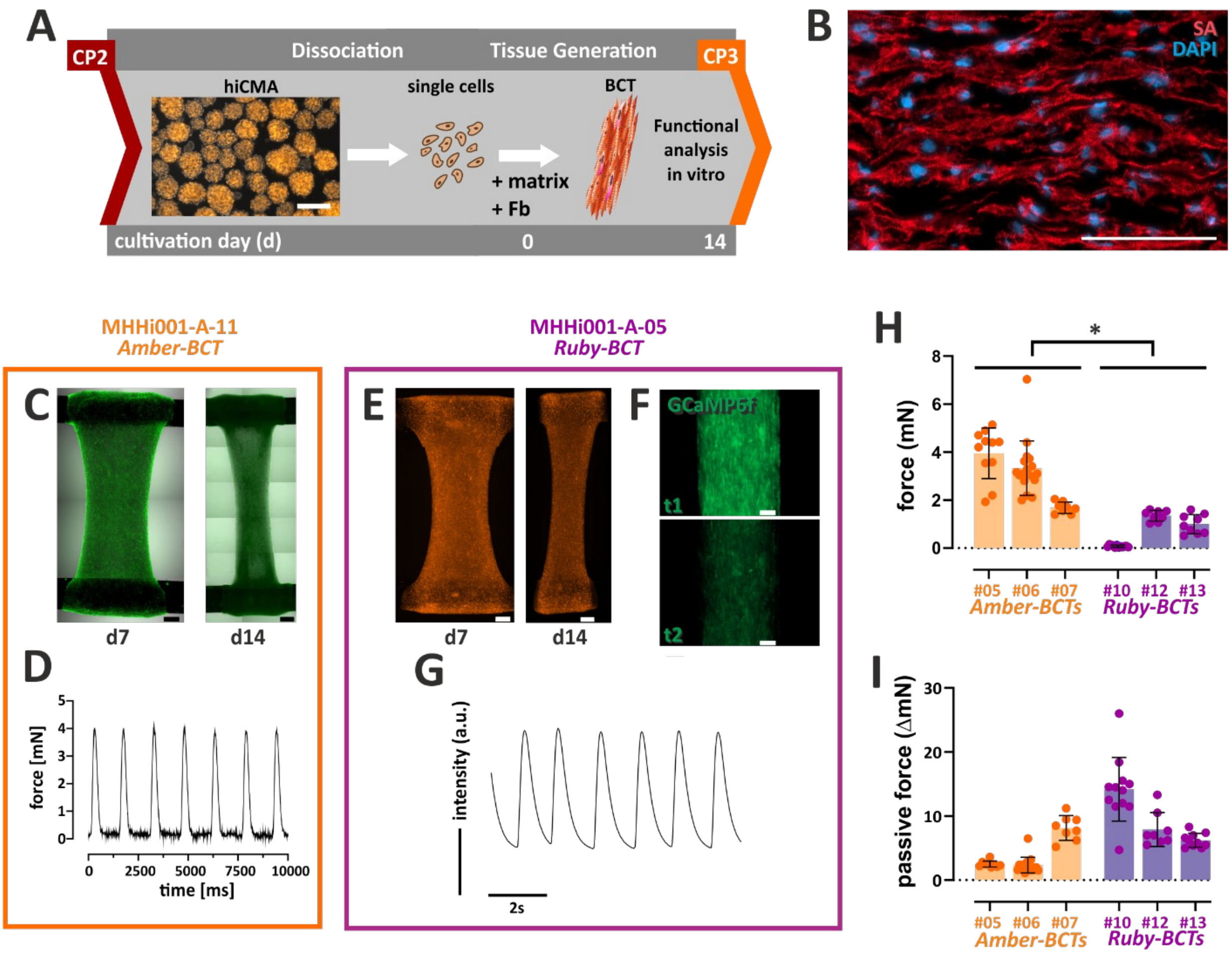


**Figure S1: In vitro formation of homogenous bioartificial cardiac tissue confirms plasticity and functionality of hiCMAs used for transplantation**

**A)** Bioartificial cardiac tissue (BCT) produced from batches of hiCMAs at the time point of transplantation (CP2) was used to assess the ability of the hiCMAs from the two cell lines MHHi0001-A-11 ‘*Amber*’ (**B-D**), or MHHi0001-A-5 ‘*Ruby*’ [4]) (**E-G**) to form structured myocardium in vitro. After 14 d of cultivation (CP3), BCTs were used for functional analysis of CM contractility of individual batches (**H, I**). **B)** Immunofluorescence (IF) staining of cryosections of d14 BCT for α-sarcomeric actinin (SA, red) showed elongated and aligned CMs and a cross-striated staining pattern that reflects sarcomeric organization. Nuclei were stained with DAPI (blue), scale bar 100 µm. **C)** BCTs from MHHi0001-A-11 ‘*Amber*’ (expressing the Venus(nucmem) transgene) show viable CMs with tissue remodelling on d7 and d14; scale bars 500 µm. **D)** Exemplary force measurement data from a spontaneously beating Amber-BCT acquired with force transducers in a custom-made bioreactor system. **E)** BCTs from MHHi0001-A-5 ‘*Ruby*’ (expressing the RedStar(nucmem) and GCamP6f transgenes) show viable CMs with tissue remodeling on d7 and d14; scale bars 500 µm. **F)** Expression of the calcium sensor GCamP6f in Ruby-BCTs allowed microscopic monitoring of intracellular calcium in BCTs (still images from video-optical analysis, scale bar 200 µm). **G)** Video-optical analysis of calcium-dependent fluorescence oscillations confirm BCT contractility (a.u. arbitrary units). **H)** On d14, maximum active forces of BCTs from individual hiCMA batches show large variations of contractility in vitro with significantly higher forces for Amber-BCTs vs. Ruby-BCTs (3.2 ± 1.3 mN vs. 0.7 ± 0.6 mN; Mean ± SD for n=8-17 BCTs from 3 experiments each; $p<0.05$ for nested t-test). **I)** Passive forces indicating tissue stiffness tended to be higher in Ruby-BCTs than in Amber-BCTs with values up to 14.2 ± 5.0 mN (Mean ± SD, n=12 BCTs) for the least contractile Ruby-BCTs from hiCMA batch #10.

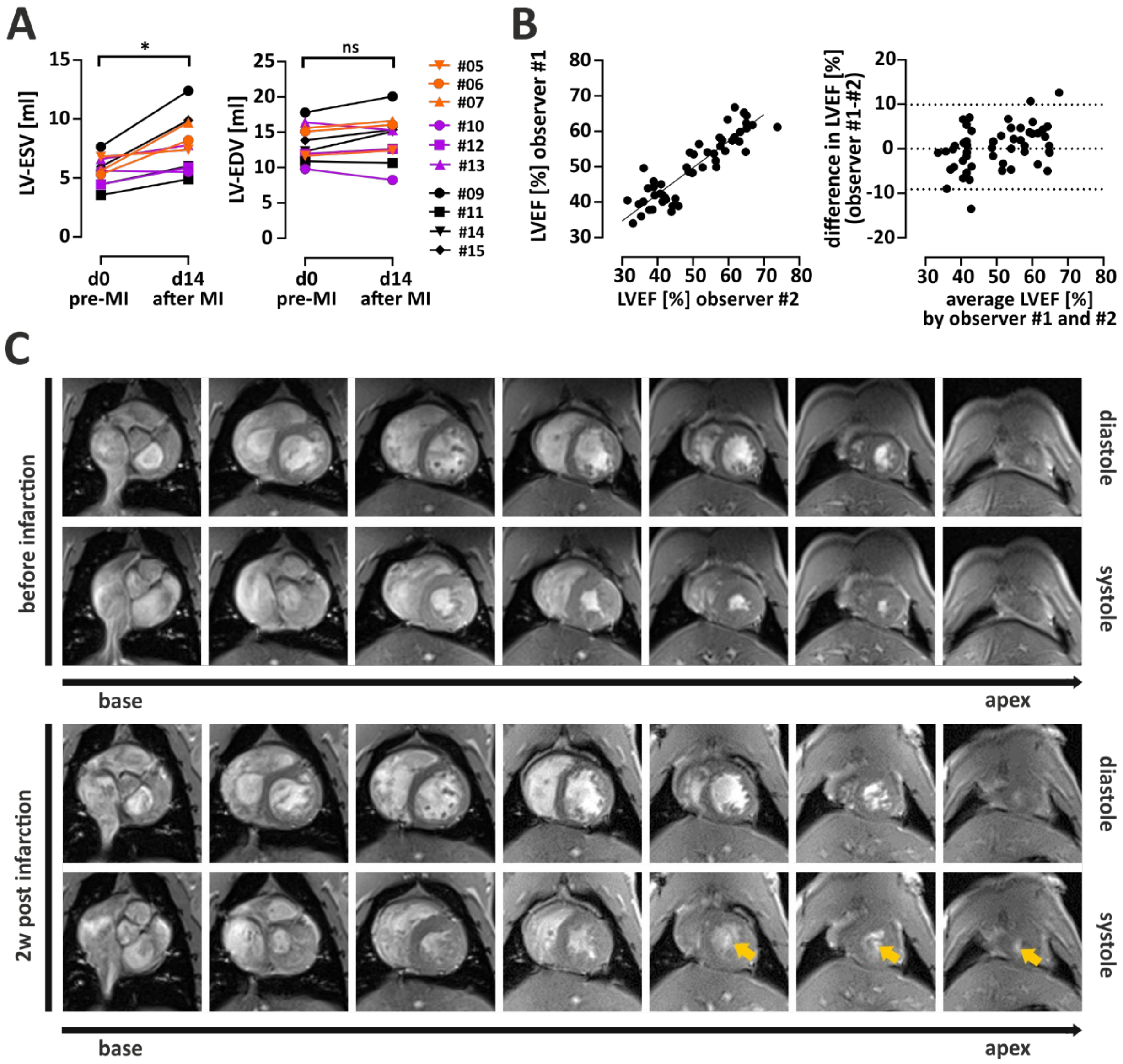


**Figure S2: Cardiac MRI assessment of the infarcted myocardium showed impaired function and infarct localization**

**A)** Cardiac MRI was performed to measure left ventricular end-systolic volume (LV-ESV, left panel) and left ventricular end-diastolic volume (LV-EDV, right panel) pre-MI and on d14 after MI for individual animals. The increase of LV-ESV after MI was significant (7.8 ± 2.4 ml vs. 5.6 ± 1.2 ml before MI; $p<0.05$ for n=10, paired-t-test), while LV-EDV did not change significantly (see table 1 for individual data). The infarcted animals were assigned to the treatment groups A and B1 (treatment with hiCMA '*Amber*', orange symbols), group B2 (treatment with hiCMA '*Ruby*', purple symbols), and group C (sham, black). **B)** Good inter-observer agreement was achieved for the left ventricular ejection fraction (LV-EF) calculated from LV-ESV and LV-EDV; left panel), the bias (difference between the means) was only 0.4 and the intra-class correlation coefficient (ICC) 0.89. Bland-Altman plot shows the inter-observer variability with 95% limits of agreement between -9.086 and 9.890. **C)** Short-axis images of selected slices of monkey #09 before and 2 weeks after myocardial infarction (MI) from base to apex. A significantly larger systolic volume in comparison to the non-infracted heart is clearly visible on the apical slices 2 weeks after MI (see arrows) and shows loss of systolic contractility.

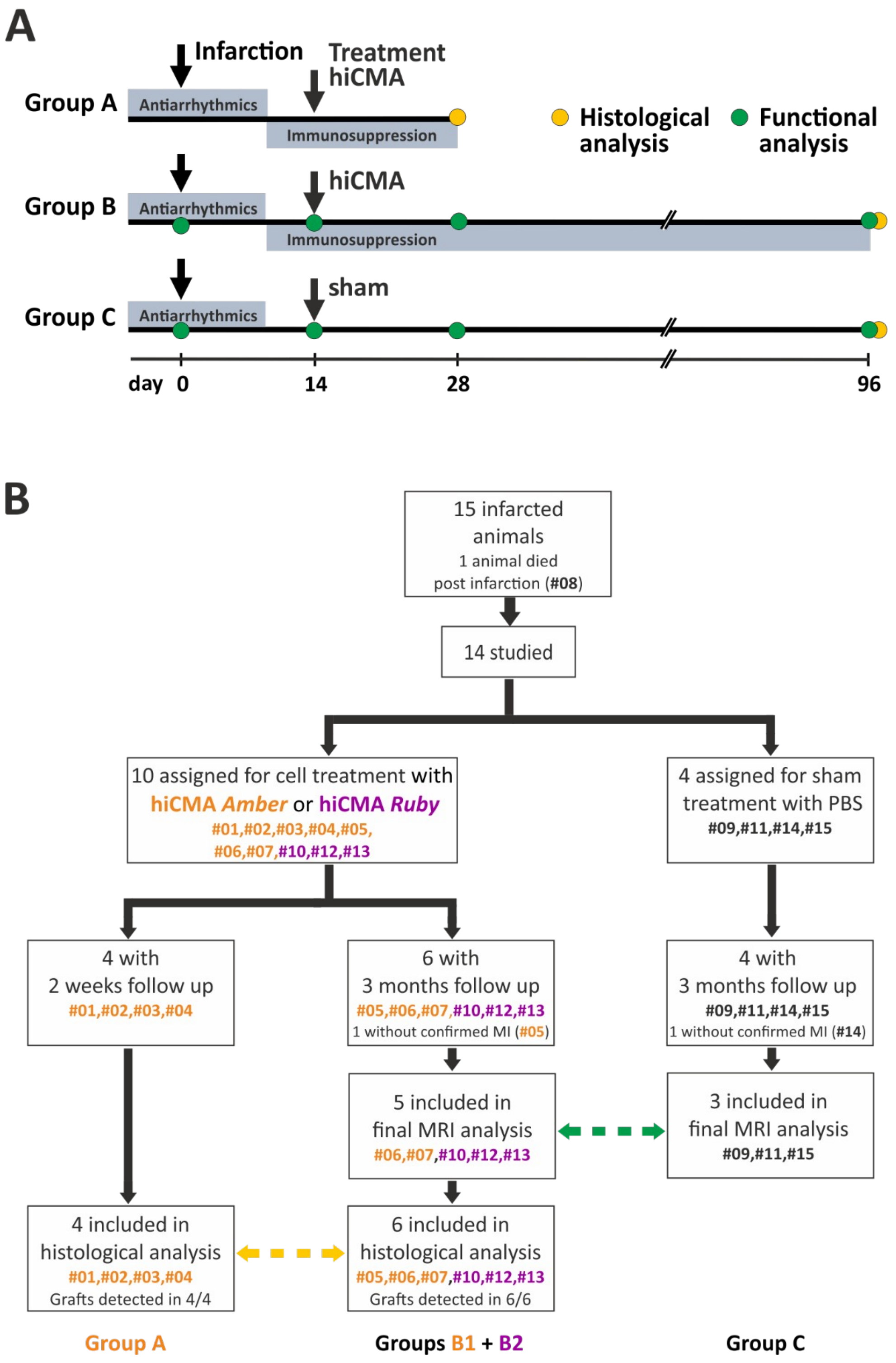


**Figure S3: Animal study design**

**A)** Time line for interventions and analyses. The simplified scheme shows i) time points of major interventions, ii) timing of medication with anti-arrhythmics and immunosuppressants, and iii) time points for functional analysis by MRI and histological analysis for the different experimental groups A, B and C. **B)** Flow diagram for animal assignment to treatment groups and inclusion in MRI and histology analysis. The figure depicts the experimental flow, mortality at each stage of the study, assignment of individual animals into experimental groups, and final group sizes for MRI and histology analyses.

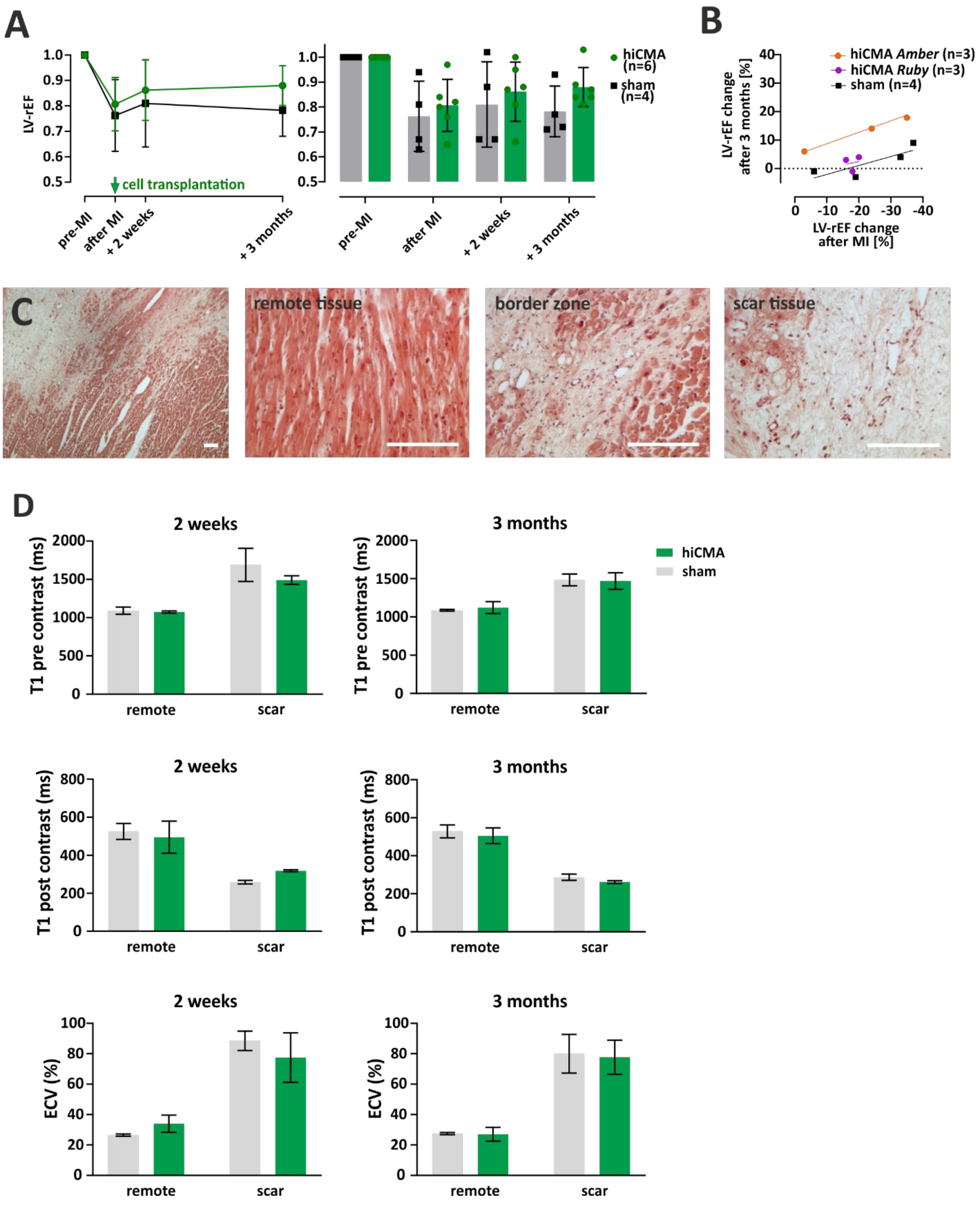


**Figure S4: Extended cardiac MRI data assessment and scar tissue characterization after up to 3 months**

**A)** Due to the large variance of pre-MI LVEF in cardiac MRI measurements, LV-rEF values normalized to pre-MI values were used to assess the outcome of individual hiCMA-treated animals. For the combined assessment of treatment groups B1+B2 (green symbols, n=6; #05, #06, #07, #10, #12, #13) compared to the sham control group C (black symbols, n= 4; #09, #11, #14, #15), a trend for increased LVEF was observed after 3 months, but did not reach statistical significance (0.88 ± 0.08 vs. 0.78 ± 0.10; p = 0.5; left panel). The right panel displays the large variation of values within groups, most probably preventing statistical significance. Notably, the two animals with very small LV-EF changes after MI (animals #05 and #14, Fig. 2C) could be identified as outliers after 2 weeks and 3 months as well, because they did not show major changes after treatment or sham. **B)** Re-analysis of the MRI data showed that the observed functional improvement (LV-rEF change after 3 months) appears to be correlated to the initial decrease in function (LV-rEF change 2 weeks after MI) and is clearly dependent on the treatment group. A slight recovery was observed for sham treated animals after large impairment (>30% LV-rEF reduction), this was clearly outperformed after treatment with hiCMA from MHHi0001-A-11 '*Amber*' (as can be seen by the difference in elevation of the linear regression), but not after treatment with hiCMA from MHHi0001-A-5 'Ruby'. **C)** Microscopic pictures after HE staining confirmed the presence of infarcted myocardium and scar tissue formation in an untreated animal (#09) after 3 months. Scale bars 200 µm. **D)** Myocardial tissue characteristics and fibrotic changes were assessed by cardiac MRI, i.e. T1-relaxation times of scar and remote myocardium prior and 15 minutes after contrast agent administration. The scar regions showed an increase in the pre-contrast (top) and a decrease in the post-contrast (middle) T1-relaxation time compared to the remote region as expected after tissue injury. No significant difference could be observed between the treated and untreated animals, neither 2 weeks nor 3 months after treatment. The scar tissue showed an increased extracellular volume (ECV) independent of the treatment (bottom), demonstrating the presence of fibrotic tissue in all animals, however, the size of the scarred area was not assessed.

### Supplemental videos

**Video SV1: hiCMAs derived from the reporter iPSC line "*Amber*"**
Light microscopy video representative of contractile CM-aggregates, sampled and assessed at check point 2 (CP2). Note the well-defined smooth outer appearance of each aggregate and the lack of single cells. Scale bar (lower right corner) represents 200 µm.

**Video SV2: hiCMAs derived from the reporter iPSC line "*Ruby*"**
Fluorescence microscopy video representative of contractile CM-aggregates, sampled and assessed at CP2. The expected flashing of individual aggregates over time is evoked by calcium transients visualised by the reporter line specific transgenic calcium sensor GCaMP6f. Scale bar (lower right corner) represents 200 µm.

### Supplemental References


1. Manstein F, Ullmann K, Kropp C, Halloin C, Triebert W, Franke A, Farr CM, Sahabian A, Haase A, Breitkreuz Y, Peitz M, Brüstle O, Kalies S, Martin U, Olmer R, Zweigerdt R. High density bioprocessing of human pluripotent stem cells by metabolic control and in silico modeling. Stem Cells Transl Med 2021.
2. Halloin C, Schwanke K, Lobel W, Franke A, Szepes M, Biswanath S, Wunderlich S, Merkert S, Weber N, Osten F, de la Roche J, Polten F, Wollert K, Kraft T, Fischer M, Martin U, Gruh I, Kempf H, Zweigerdt R. Continuous WNT Control Enables Advanced hPSC Cardiac Processing and Prognostic Surface Marker Identification in Chemically Defined Suspension Culture. Stem Cell Reports 2019;13(2):366-379.
3. Nagai T, Ibata K, Park ES, Kubota M, Mikoshiba K, Miyawaki A. A variant of yellow fluorescent protein with fast and efficient maturation for cell-biological applications. Nat Biotechnol 2002;**20**(1):87-90.
4. Haase A, Kohrn T, Fricke V, Ricci Signorini ME, Witte M, Göhring G, Gruh I, Martin U. Establishment of MHHi001-A-5, a GCaMP6f and RedStar dual reporter human iPSC line for in vitro and in vivo characterization and in situ tracing of iPSC derivatives. Stem Cell Res 2021;**52**:102206.
5. Palecek J, Zweigerdt R, Olmer R, Martin U, Kirschning A, Drager G. A practical synthesis of Rho-Kinase inhibitor Y-27632 and fluoro derivatives and their evaluation in human pluripotent stem cells. Org Biomol Chem 2011;**9**(15):5503-10.
6. Kensah G, Roa Lara A, Dahlmann J, Zweigerdt R, Schwanke K, Hegermann J, Skvorc D, Gawol A, Azizian A, Wagner S, Maier LS, Krause A, Drager G, Ochs M, Haverich A, Gruh I, Martin U. Murine and human pluripotent stem cell-derived cardiac bodies form contractile myocardial tissue in vitro. Eur Heart J 2013;**34**(15):1134-46.
7. Tian Y, Martinez MM, Pappas D. Fluorescence correlation spectroscopy: a review of biochemical and microfluidic applications. Appl Spectrosc 2011;**65**(4):115a-124a.
8. Liu YW, Chen B, Yang X, Fugate JA, Kalucki FA, Futakuchi-Tsuchida A, Couture L, Vogel KW, Astley CA, Baldessari A, Ogle J, Don CW, Steinberg ZL, Seslar SP, Tuck SA, Tsuchida H, Naumova AV, Dupras SK, Lyu MS, Lee J, Hailey DW, Reinecke H, Pabon L, Fryer BH, MacLellan WR, Thies RS, Murry CE. Human embryonic stem cell-derived cardiomyocytes restore function in infarcted hearts of non-human primates. Nat Biotechnol 2018;**36**(7):597-605.
9. Moussavi A, Mißbach S, Serrano Ferrel C, Ghasemipour H, Kötz K, Drummer C, Behr R, Zimmermann WH, Boretius S. Comparison of cine and real-time cardiac MRI in rhesus macaques. Sci Rep 2021;**11**(1):10713.
10. Heiberg E, Sjögren J, Ugander M, Carlsson M, Engblom H, Arheden H. Design and validation of Segment--freely available software for cardiovascular image analysis. BMC Med Imaging 2010;**10**:1.
11. Wang X, Joseph AA, Kalentev O, Merboldt KD, Voit D, Roeloffs VB, van Zalk M, Frahm J. High-resolution myocardial T(1) mapping using single-shot inversion recovery fast low-angle shot MRI with radial undersampling and iterative reconstruction. Br J Radiol 2016;**89**(1068):20160255.
